# Multiple, independent, common variants overlapping known and putative gut enhancers at *RET*, *SEMA3* and *NRG1* underlie Hirschsprung disease risk in European ancestry subjects

**DOI:** 10.1101/2020.06.07.138719

**Authors:** Ashish Kapoor, Priyanka Nandakumar, Dallas R. Auer, Maria X. Sosa, Holly Ross, Juli Bollinger, Jia Yan, Courtney Berrios, The Hirschsprung Disease Research Collaborative (HDRC), Aravinda Chakravarti

## Abstract

**Purpose:** Hirschsprung disease (HSCR) is a developmental disorder of the enteric nervous system (ENS) characterized by congenital aganglionosis, and where individual cases harbor coding risk variants in ENS genes. Low-penetrance, common, noncoding variants at RET, SEMA3 and NRG1 loci have been associated in HSCR as well, implicating variable gene expression mediated by cis-regulatory element (CRE) variants as the causal mechanism. However, the extent and combinatorial effects of all putative CRE variants within and across these loci on HSCR is unknown.

**Methods:** Using 583 HSCR subjects, one of the largest samples of European ancestry studied, and genotyping 56 tag variants we evaluated association of all common variants overlapping putative gut CREs and fine-mapped variants at RET, SEMA3 and NRG1.

**Results:** We demonstrate that 28 and 8 tag variants, several of which are independent, overlapping putative-enhancers at the RET and SEMA3 loci, respectively, and, 2 fine-mapped tag variants at NRG1 locus, are associated with HSCR. We demonstrate that risk increases with increasing risk allele dosage from multiple variants within and across these loci and varies >25- fold.

**Conclusion:** Gene regulatory networks in HSCR-relevant cell types quantify the total burden of risk alleles through sensing reduced gene expression of multiple genes on disease.

## Introduction

Hirschsprung disease (HSCR) is a developmental disorder of the enteric nervous system (ENS), characterized by loss of neuronal ganglia in the myenteric and submucosal plexuses of the gut.^1^ Failure of the precursor enteric neural crest cells to proliferate, differentiate, migrate and/or colonize the gastrointestinal tract leads to HSCR, the most common (15 per 100,000 live births^2^) cause of functional intestinal obstruction in neonates and infants.^3^ The ensuing aganglionosis is caudal to rostral and based on the extent of aganglionosis, patients are classified into short-segment HSCR (S-HSCR; aganglionosis limited up to the upper sigmoid colon, ∼80% of cases), long-segment HSCR (L-HSCR; aganglionosis up to splenic flexure and beyond, ∼15% of cases) or total colonic aganglionosis (TCA; aganglionosis of the entire large intestine, ∼5% of cases).^1^ Nearly 80% of cases have aganglionosis only; in the remainder, HSCR co-occurs with multiple congenital anomalies, specific syndromes and/or chromosomal variants.^1,4^ The critical features of HSCR are its high heritability (>80%), complex inheritance pattern, sex bias (4:1 affected male: female) and high recurrence risk to siblings and other relatives, influenced by sex, segment length of aganglionosis, familiality and the presence of additional anomalies.^2^

Genetic studies in HSCR have uncovered rare, high-penetrance, coding variants in 14 genes^4–6^ and common, low-penetrance, noncoding variants near *RET*^6–8^, *NRG1*^9^ and the *SEMA3* gene cluster.^6^ The pathogenic coding alleles at these critical ENS genes collectively lead to a population attributable risk (PAR) of ∼18.2%.^10^ In contrast, the polymorphic noncoding variants at *RET* and *SEMA3* lead to a much larger PAR of ∼37.7%.^10^ Although these noncoding variants at *RET* and *SEMA3* individually confer low-to-moderate risk (odds ratio, OR: 1.6-3.9), collectively, they contribute to a 30-fold *risk variation* depending on risk allele dosage.^11^ While variable *RET* expression mediated by noncoding regulatory variants is the mechanism at *RET*, the causal molecular basis of associations at *NRG1* and *SEMA3* remain unexplored. Functional studies at the *RET* locus, have proven that three risk variants lead to reduced *RET* expression by disrupting SOX10, GATA2 and RARB binding at three functionally distinct but synergistically acting *RET* gut enhancers or *cis*-regulatory elements (CREs).^8,12^ Although these three *RET* CRE variants are genetically independent (based on linkage disequilibrium (LD) in controls) they have epistatic effects on risk.^11,12^

Given the widespread nature of multiple putative CREs for human genes, uncovered by the ENCODE^13^ and NIH RoadMap Epigenomics^14^ Projects, and supported by studies in model systems,^15^ along with the extensive common noncoding variation in humans,^16,17^ it is *a priori* likely that *multiple CRE variants* at any given disease-associated locus can affect gene expression and thereby modulate disease risk, as shown for *RET*.^12^ Thus, multiple CRE variants affect a target gene’s expression in *cis* and suggest haplotype effects independent of the LD between risk variants.^12^ Finally, if larger numbers of variants lead to more extreme gene expression changes, and therefore higher risk, then affected individuals will be enriched for multiple risk variants. Indeed, there are considerable hints, from prior segregation^2^, linkage^18,19^, and association^11^ studies that multiple risk variants define HSCR in *individuals*. In other words, mechanistically, an association locus is likely to have multiple causal CRE variants modulating gene expression and disease risk.^20^

In this study, we have attempted to identify multiple causal variants by evaluating association of *all* common noncoding variants overlapping HSCR-relevant putative enhancers at the *RET* and *SEMA3* loci with HSCR risk, in a large sample of European ancestry cases. We also include all fine-mapped variants at the *NRG1* locus^21^, associated with HSCR risk in Asian ancestry subjects, to reassess whether this association also exists in this larger European ancestry sample. We indeed demonstrate that multiple, independent, putative-enhancer overlapping variants at the *RET* and *SEMA3* loci, and fine-mapped variants at *NRG1* locus, associate with HSCR risk. We also demonstrate that this risk increases as a logistic function with increasing risk allele dosage from multiple variants across these loci, suggesting that enteric neurons can quantitatively assess gene expression status across these, and presumably other, genes through gene regulatory networks (GRN).

## Materials and Methods

### Patient samples

We analyzed HSCR patients and their family members ascertained from two sources: ongoing family studies from this (Chakravarti) laboratory (AC-HSCR) and by the Hirschsprung Disease Research Collaborative (HDRC; Appendix S1). The AC-HSCR collection includes participants recruited from referrals by practicing physicians, genetic counselors, family members and from self-referral through a study website and support group postings (*n*=461 families). The HDRC is a multi-institutional study that recruits participants from newly diagnosed cases from pediatric surgery and clinical centers at HDRC institutions (*n*=276 families). The majority of AC-HSCR and HDRC participants are of self-described European ancestry.

### Ethics statement

All individuals were ascertained with written informed consent approved by the Institutional Review Board of the enrolling institution. This study is based on samples approved by the Institutional Review Board of the Johns Hopkins School of Medicine (March 31, 2018).

### Variant selection

For the *RET* (Figure S1) and *SEMA3* (Figure S2) loci, we defined the target regions based on HSCR genome-wide association study (GWAS) signals^6^ and flanking recombination hotspots from HapMap^16^. Within these regions we selected common variants (minor allele frequency (MAF) >10%) observed in 1000 Genomes^17^ non-Finnish European (NFE) ancestry (*n*=404) that overlapped putative gut enhancer marks from public epigenomic databases,^13,14^ as described earlier^12^ (Dataset S1). Due to lack of prior evidence for association at *NRG1* locus in European ancestry HSCR subjects, we didn’t perform the extensive association screen but selected all 17 fine-mapped variants reported to be associated with HSCR in Asian ancestry subjects (Dataset S1).^21^ All variants selected at a locus fall within the same topologically-associated domain as the underlying risk gene (*RET*, *SEMA3D* and *NRG1*).^22^ Using 1000 Genomes^17^ NFE ancestry genotype data and the software Tagger^23^, 42, 11 and 7 tag variants at the *RET*, *SEMA3* and *NRG1* loci, respectively were selected with a LD *r*^2^ cutoff of 0.5. We added the top two GWAS hits at *SEMA3* locus (rs11766001 and rs12707682)^6^ for a total of 62 variants that underwent multiplex genotyping assay design (Dataset S1).

### Multiplex genotyping assay

All steps of genotyping, including the design of custom multiplexed assays using iPLEX chemistry, were performed following the manufacturer’s recommendations (Agena Bioscience, San Diego, CA, USA). Variants that failed *in silico* assay design or failed in genotyping of reference DNA samples were replaced with a best proxy variant from the filtered sets of variants when available; if not, we used the best replacement from the larger sets of common variants (based on LD) to generate two multiplexed assay pools for 61 tag variants (Dataset S1).

### Variant genotyping quality control (QC)

Genotyping of 2,205 unique DNA samples was carried out for each multiplexed pool. Post sample and variant QC (Supplementary Methods, Dataset S1, Table S1, S2) we combined variants across the two pools (31 and 25 variants in pools #1 and #2, respectively) to generate genotype data for 56 variants in 1,959 samples from 724 families.

**Table 1:** Distribution of HSCR probands from the AC-HSCR and HDRC studies by sex, segment length of aganglionosis and familiality.

| Category | Combined Count (%) | AC-HSCR Count (%) | HDRC Count (%) | Test of frequency difference |  |
| --- | --- | --- | --- | --- | --- |
| All | 583 (100) | 375 (100) | 208 (100) |  |  |
| | | | | $\chi^2$ | *P* |
| **Sex** |  |  |  |  |  |
| Male | 425 (72.9) | 267 (71.2) | 158 (76.0) | 1.54<sup>a</sup> | 2.15×10<sup>-1</sup> |
| Female | 158 (27.1) | 108 (28.8) | 50 (24.0) |  |  |
| **Segment length** |  |  |  |  |  |
| Short | 274 (47.0) | 162 (43.2) | 112 (53.8) | 7.22<sup>b</sup> | 7.23×10<sup>-3</sup> |
| Long | 57 (9.8) | 42 (11.2) | 15 (7.2) |  |  |
| TCA | 91 (15.6) | 65 (17.3) | 26 (12.5) |  |  |
| Indeterminate | 161 (27.6) | 106 (28.3) | 55 (26.4) |  |  |
| **Familiarity** |  |  |  |  |  |
| Simplex | 425 (72.9) | 249 (66.4) | 176 (84.6) | 22.47<sup>c</sup> | 2.14×10<sup>-6</sup> |
| Multiplex | 158 (27.1) | 126 (33.6) | 32 (15.4) |  |  |

<sup>a</sup>Comparison between males and females, <sup>b</sup>short and long+TCA cases, and <sup>c</sup>simplex and multiplex cases, respectively.

Genotypes at each variant from all unrelated HSCR probands (*n*=583) were tested for Hardy-Weinberg Equilibrium (HWE): none were significant (*P*<1.0×10^-6^) except rs2435357 (*P*=2.69×10^-19^; Table S3), which was expected owing to its high population-level association with HSCR.^7,8^ None of the HWE tests using genotypes from 1000 Genomes^17^ NFE ancestry subjects were significant (Table S3).

**Table 3:** Odds ratio and risk of HSCR as a function of the number of risk-increasing variants at *RET*, *SEMA3* and *NRG1*.

| Risk allele counts | Number (%) of cases | Number (%) of controls | Odds ratio (95% CI) | *P* | Penetrance (x10<sup>5</sup>) |
| --- | --- | --- | --- | --- | --- |
| 19-29 | 25 (5.0) | 82 (20.3) | **0.21** (0.13-0.33) | 5.64×10<sup>-11</sup> | 3.7 |
| 30-33 | 44 (8.9) | 89 (22.0) | **0.34** (0.23-0.51) | 7.32×10<sup>-8</sup> | 6.0 |
| 34-37 | 66 (13.3) | 88 (21.8) | **0.55** (0.39-0.78) | 8.28×10<sup>-4</sup> | 9.1 |
| 38-42 | 94 (18.9) | 73 (18.1) | 1.06 (0.75-1.48) | 7.46×10<sup>-1</sup> | 15.7 |
| 43-66 | 268 (53.9) | 72 (17.8) | **5.40** (3.96-7.36) | 1.54×10<sup>-26</sup> | 45.4 |

Overall $\chi^2=156.43$ ; $P=8.52\times 10^{-33}$
The numbers of cases and controls classified by the number of risk alleles at *RET*, *SEMA3* and *NRG1* significant variants, odds ratio with 95% confidence interval (CI), statistical significance of association (*P*) and estimated population incidence (penetrance) are provided. Statistically significant values are in bold. The penetrance value shown is the expected number of cases in 100,000 live births of that class assuming a total population incidence of 15 cases per 100,000 live births.

### Statistical genetic analyses

For control allele frequencies, we used publicly available data from the Genome Aggregation Database (gnomAD)^24^, based on whole-genome sequencing of NFE ancestry subjects. All HSCR cases with complete information on sex, segment length of aganglionosis and familiality, were classified into 8 risk categories, scored by known HSCR risk factors (score of 0 each for male, short segment and simplex versus a score of 1 each for female, long segment/TCA and multiplex), and eventually summed into risk scores between 0-3 (Table S4)^25^. Population-level disease penetrance for each risk allele bin was estimated using Bayes’ theorem with the observed background control frequency and a disease incidence of 15 per 100,000 live births.^8^ Polygenic risk scores (PRS) were calculated as the sum of risk alleles at HSCR-associated variants weighted by log10-transformed OR from case-control tests (AC-HSCR and HDRC combined cases vs NFE controls). Statistical significance thresholds for single variant-based tests were adjusted for multiple tests using the Bonferroni correction for 56 tests at *P*<8.9×10^-4^. All *P*-values reported are two-tailed.

Additional details of Methods are in Supplementary Material.

## Results

### HSCR subtypes differ between cohorts

We used two HSCR sample cohorts in this study: AC-HSCR and HDRC. AC-HSCR, collected through this laboratory, is likely to over-sample high-risk families while HDRC collected at clinical centers at first surgical encounter is likely to be representative of HSCR in the general population (Appendix S1). To assess if ascertainment differences can influence our analysis (see below), we compared the proportions of HSCR subtypes by sex, segment length (S- HSCR versus L-HSCR/TCA) and familiality in probands across the two cohorts (Table 1).

Ignoring samples with unknown status, there was no significant difference between the cohorts for sex (71% vs. 76% males in AC-HSCR and HDRC, respectively; *P*=0.22). However, HDRC was significantly enriched for S-HSCR and simplex cases as compared to AC-HSCR (73% vs. 60% S-HSCR, *P*=0.007; 85% vs. 66% simplex, *P*=2.1×10^-6^ in HDRC and AC-HSCR, respectively). This confirms the expectation that AC-HSCR was enriched for high-risk families, i.e. those with L-HSCR/TCA and with affected relatives. These data also suggest that, in unselected HSCR cases the frequency of males, L-HSCR/TCA and positive family history are 76%, 27% and 15%, respectively. We also classified each proband into 8 risk categories based on known HSCR risk factors,^25^ and four risk classes (scores 0-3; Table S4). Comparisons of risk scores 0, 1 and 2+3 (combined due to small sample sizes) between the two cohorts showed a highly significant difference (*P*=3.3×10^-4^), with groups 0 and 2+3 being more abundant in HDRC and AC-HSCR, respectively (Table S5). Thus, the two samples may reveal different genotype-phenotype correlations.^8^

### Widespread genetic association of RET, SEMA3 and NRG1 variants with HSCR

Previous studies have established that common noncoding variants at *RET*, *SEMA3* and *NRG1* are significantly associated with HSCR risk.^6–9^ However, these studies were from gene discovery centers who may have over-sampled probands from high-risk families. This is relevant because our first discovered genetic association had differing phenotypic features depending on HSCR risk factors.^7,8,11^ Our goal here, for the *RET* and *SEMA3* loci, was therefore to assess the numbers and degree of genetic associations at functionally relevant CREs, in HSCR cases including randomly ascertained new samples (HDRC), and to test for the presence of independent signals within each locus. We also wished to reassess the HSCR association^9^ at *NRG1*. We genotyped a total of 2,205 subjects across 737 families from AC-HSCR and HDRC for 61 tag variants (MAF>10%), which after QC (Dataset S1, Tables S1-S3) reduced to genotype data from 56 tag variants in 1,959 subjects (701 affected) from 724 families (583 probands).

From these, data from 583 independent HSCR cases were used for association analyses (Table 1). Of the 375 AC-HSCR independent cases, 320 have been evaluated earlier for association with limited set of risk variants at *RET*, *SEMA3* and *NRG1*.^11,12^

We used population-based case-control analysis and compared allele frequencies between HSCR cases and NFE ancestry controls (Table 2). At *RET*, 27 of 37 tag variants showed significant association with HSCR, including three previously published independent associations at rs2435357 (60% vs. 27%, OR=4.0, *P*=1.1×10^-123^)^7,8,11^, rs2506030 (56% vs. 39% at rs788261, OR=2.0, *P*=2.8×10^-30^, *r^2^*=0.98 with 2506030)^6,11^ and rs7069590 (84% vs. 77%, OR=1.5, *P*=1.8×10^-7^).^12^ Among all *RET* variants evaluated, the most significant association remained at rs2435357. The ORs observed here at rs2435357, rs2506030 and rs7069590 are very similar to that previously reported in European ancestry HSCR subjects, in overlapping samples from AC-HSCR and by others (ORs range: 3.9-6.7, 1.8-2.3 and 1.5-1.7 for rs2435357, rs2506030 and rs7069590, respectively; Table S6). At *SEMA3*, 8 of 12 tag variants showed significant association with HSCR, including two previously published GWAS hits at rs12707682 (31% vs. 23%, OR=1.6, *P*=2.6×10^-11^) and rs11766001 (19% vs. 14%, OR=1.5, *P*=1.3×10^-6^).^6,11^ The ORs observed here at rs12707682 and rs11766001 are also very similar to that reported in European ancestry HSCR subjects, in overlapping samples from AC-HSCR and by others (ORs range: 1.3-1.8 and 1.4-2.3 for rs12707682 and rs11766001, respectively; Table S6). The most significant association at *SEMA3* was, however, at rs1228877 (45% vs. 33%, OR=1.6, *P*=2.6×10^-16^). At *NRG1* locus, 2 variants showed significant association with HSCR, including one of the Asian ancestry HSCR GWAS hits at rs16879552 (5% vs. 3%, OR=1.9, *P*=2.7×10^-6^)^9^, a new result in European ancestry HSCR subjects.^6,11,26^ This new observation, previously not significant (*P*=0.4) in overlapping samples from AC-HSCR^11^ (Table S6), is driven by the HDRC samples (see below). The most significant association at *NRG1* was, however, at rs16879576 (5% vs. 2%, OR=2.3, *P*=7.2×10^-9^), also driven by the HDRC samples (see below). We also calculated parent to child transmission rates for all tag variants in 314 HSCR trios and observed over-transmission of risk alleles at 31 variants (*P*<0.05) (Table S7).

**Table 2:** Case-control association tests for tag variants at *SEMA3*, *NRG1* and *RET* in HSCR.

| Gene Locus | Variant | Coded allele | Allele frequency (case/control) | Odds ratio (95% CI) | *P* |
| --- | --- | --- | --- | --- | --- |
| *SEMA3* | rs10266471 | C | 0.83/0.78 | 1.37 (1.17-1.60) | **9.98×10<sup>-5</sup>** |
| *SEMA3* | rs1228877 | G | 0.45/0.33 | 1.65 (1.46-1.86) | **2.60×10<sup>-16</sup>** |
| *SEMA3* | rs55737059 | A | 0.90/0.89 | 1.04 (0.85-1.26) | 7.26×10<sup>-1</sup> |
| *SEMA3* | rs1228937 | C | 0.26/0.20 | 1.44 (1.25-1.65) | **1.95×10<sup>-7</sup>** |
| *SEMA3* | rs11766001 | C | 0.19/0.14 | 1.46 (1.25-1.71) | **1.31×10<sup>-6</sup>** |
| *SEMA3* | rs2190050 | T | 0.22/0.19 | 1.18 (1.02-1.36) | 2.80×10<sup>-2</sup> |
| *SEMA3* | rs35092834 | T | 0.73/0.66 | 1.44 (1.26-1.64) | **1.27×10<sup>-7</sup>** |
| *SEMA3* | rs12707682 | C | 0.31/0.23 | 1.56 (1.37-1.77) | **2.57×10<sup>-11</sup>** |
| *SEMA3* | rs7811476 | G | 0.32/0.30 | 1.06 (0.93-1.21) | 3.65×10<sup>-1</sup> |
| *SEMA3* | rs6468008 | G | 0.58/0.51 | 1.34 (1.19-1.52) | **1.49×10<sup>-6</sup>** |
| *SEMA3* | rs17560655 | C | 0.18/0.12 | 1.59 (1.36-1.86) | **6.96×10<sup>-9</sup>** |
| *SEMA3* | rs6974210 | G | 0.87/0.86 | 1.04 (0.87-1.24) | 6.69×10<sup>-1</sup> |
| *NRG1* | rs16879425 | C | 0.12/0.09 | 1.28 (1.06-1.54) | 9.53×10<sup>-3</sup> |
| *NRG1* | rs10954845 | G | 0.24/0.20 | 1.26 (1.10-1.45) | 1.20×10<sup>-3</sup> |
| *NRG1* | rs4422736 | T | 0.07/0.05 | 1.34 (1.06-1.69) | 1.39×10<sup>-2</sup> |
| *NRG1* | rs7826312 | C | 0.59/0.58 | 1.03 (0.91-1.16) | 6.46×10<sup>-1</sup> |
| *NRG1* | rs16879552 | T | 0.05/0.03 | 1.90 (1.45-2.50) | **2.66×10<sup>-6</sup>** |
| *NRG1* | rs7835688 | C | 0.50/0.47 | 1.14 (1.01-1.28) | 3.72×10<sup>-2</sup> |
| *NRG1* | rs16879576 | A | 0.05/0.02 | 2.34 (1.74-3.14) | **7.17×10<sup>-9</sup>** |
| *RET* | rs11239767 | T | 0.82/0.81 | 1.07 (0.92-1.25) | 3.89×10<sup>-1</sup> |
| *RET* | rs12570505 | T | 0.89/0.89 | 1.04 (0.86-1.26) | 6.82×10<sup>-1</sup> |
| *RET* | rs3758514 | T | 0.65/0.60 | 1.20 (1.06-1.35) | 4.73×10<sup>-3</sup> |
| *RET* | rs61844837 | C | 0.94/0.91 | 1.61 (1.25-2.06) | **1.54×10<sup>-4</sup>** |
| *RET* | rs2796552 | A | 0.73/0.70 | 1.15 (1.01-1.32) | 3.81×10<sup>-2</sup> |
| *RET* | rs788243 | G | 0.57/0.49 | 1.40 (1.24-1.57) | **5.74×10<sup>-8</sup>** |
| *RET* | rs2796575 | A | 0.87/0.84 | 1.27 (1.06-1.52) | 8.29×10<sup>-3</sup> |
| *RET* | rs788274 | A | 0.27/0.13 | 2.50 (2.17-2.87) | **3.87×10<sup>-40</sup>** |
| *RET* | rs788263 | G | 0.56/0.39 | 1.99 (1.77-2.25) | **4.93×10<sup>-30</sup>** |
| *RET* | rs788261 | C | 0.56/0.39 | 2.00 (1.77-2.25) | **2.83×10<sup>-30</sup>** |
| *RET* | rs788260 | A | 0.56/0.39 | 1.99 (1.76-2.25) | **5.86×10<sup>-30</sup>** |
| *RET* | rs3004255 | T | 0.47/0.40 | 1.31 (1.17-1.48) | **7.61×10<sup>-6</sup>** |
| *RET* | rs1935646 | A | 0.29/0.26 | 1.14 (1.00-1.30) | 5.43×10<sup>-2</sup> |
| *RET* | rs11819501 | C | 0.93/0.90 | 1.36 (1.08-1.70) | 7.71×10<sup>-3</sup> |
| *RET* | rs2488279 | T | 0.28/0.20 | 1.56 (1.37-1.79) | **3.70×10<sup>-11</sup>** |
| *RET* | rs57621833 | T | 0.77/0.73 | 1.24 (1.07-1.42) | 3.12×10<sup>-3</sup> |
| *RET* | rs1547930 | G | 0.83/0.76 | 1.58 (1.35-1.85) | **1.05×10<sup>-8</sup>** |
| *RET* | rs4949062 | C | 0.73/0.67 | 1.34 (1.17-1.53) | **1.77×10<sup>-5</sup>** |
| *RET* | rs2115816 | A | 0.40/0.36 | 1.18 (1.04-1.33) | 7.78×10<sup>-3</sup> |
| *RET* | rs3026693 | A | 0.21/0.16 | 1.41 (1.21-1.63) | **6.45×10<sup>-6</sup>** |
| *RET* | rs7069590 | T | 0.84/0.77 | 1.53 (1.30-1.79) | **1.79×10<sup>-7</sup>** |
| *RET* | rs2435357 | T | 0.60/0.27 | 4.02 (3.56-4.55) | **1.13×10<sup>-123</sup>** |
| *RET* | rs12247456 | G | 0.80/0.68 | 1.84 (1.59-2.14) | **1.04×10<sup>-16</sup>** |
| *RET* | rs7393733 | G | 0.79/0.68 | 1.83 (1.58-2.12) | **2.32×10<sup>-16</sup>** |
| *RET* | rs2506024 | A | 0.80/0.60 | 2.68 (2.31-3.11) | **9.76×10<sup>-42</sup>** |
| *RET* | rs78146028 | A | 0.95/0.89 | 2.30 (1.77-3.00) | **1.55×10<sup>-10</sup>** |
| *RET* | rs12764797 | G | 0.95/0.91 | 2.05 (1.55-2.70) | **2.06×10<sup>-7</sup>** |
| *RET* | rs2505515 | G | 0.88/0.75 | 2.46 (2.05-2.95) | **8.38×10<sup>-24</sup>** |
| *RET* | rs1800860 | G | 0.74/0.68 | 1.33 (1.16-1.53) | **3.63×10<sup>-5</sup>** |
| *RET* | rs11238441 | C | 0.91/0.82 | 2.41 (1.95-2.97) | **3.21×10<sup>-17</sup>** |
| *RET* | rs2472737 | A | 0.29/0.22 | 1.40 (1.23-1.60) | **5.62×10<sup>-7</sup>** |
| *RET* | rs2505513 | C | 0.39/0.25 | 1.90 (1.67-2.15) | **1.95×10<sup>-24</sup>** |
| *RET* | rs10793423 | G | 0.52/0.44 | 1.37 (1.21-1.54) | **2.97×10<sup>-7</sup>** |
| *RET* | rs869184 | T | 0.69/0.53 | 1.93 (1.70-2.20) | **1.16×10<sup>-24</sup>** |
| *RET* | rs953999 | G | 0.80/0.68 | 1.94 (1.67-2.25) | **6.13×10<sup>-19</sup>** |
| *RET* | rs72783255 | C | 0.89/0.86 | 1.28 (1.06-1.54) | 1.07×10<sup>-2</sup> |
| *RET* | rs7074964 | A | 0.42/0.29 | 1.79 (1.59-2.03) | **3.88×10<sup>-21</sup>** |

The tag variants at each locus are listed in genomic order. The frequency of the coded allele (higher frequency in HSCR) at each variant in cases and controls, odds ratio with 95% confidence interval (CI) and the statistical significance of association (*P*) are provided. Multiple test-corrected significant *P*-values are in bold.

We calculated pairwise LD (*r*^2^) between all tag variants at each of the three loci in HSCR cases, as well as the 1000 Genomes^17^ NFE controls, to assess the underlying LD structure (Figures S3-S5) and identify associations at independent variants. Based on these results, several new and independent association signals at *RET* (Figure S3) and *SEMA3* (Figure S4) are evident. However, the two *NRG1* variants rs16879552 and rs16879576 are in moderate LD (*r*^2^=0.53/0.73 in controls/cases) (Figure S5) and may represent a singular causal association. Unlike the *RET* locus, *SEMA3* and *NRG1* loci have been largely associated with HSCR risk in European and Asian ancestry subjects, respectively. Having detected new significant associations at *NRG1* variants in European ancestry subjects here, we assessed if allele frequency differences between the two ancestries, in addition to ascertainment differences (see below), could explain these observed population-specific associations. Indeed, in general, allele frequencies for tag variants, including the GWAS hits at *SEMA3* and *NRG1* loci are quite different between NFE and East Asian gnomAD controls^24^ (Figure S6).

Given the ascertainment differences in AC-HSCR versus HDRC, we performed association analysis for all tag variants in them separately (Table S8). At *SEMA3*, the risk allele at rs12707682 was at higher frequency and led to an increased OR in HDRC (23%/29%/36% allele frequency in controls/AC-HSCR/HDRC with significant ORs of 1.4 (95% CI: 1.17-1.63) vs. 1.9 (95% CI: 1.55-2.33) in AC-HSCR and HDRC, respectively). However, within the same locus, significant risks at rs11766001 and rs1228937 were only evident in AC-HSCR (14%/21%/16% and 20%/28%/23% allele frequency in controls/AC-HSCR/HDRC with ORs of 1.7 (95% CI: 1.38-2.00) vs. 1.1 (95% CI: 0.87-1.50) and 1.6 (95% CI: 1.32-1.84) vs. 1.2 (95% CI: 0.98-1.55) in AC-HSCR and HDRC for rs11766001 and rs1228937, respectively). The confidence intervals for the above three *SEMA3* variants overlap and so the estimates are not significantly different between the two cohorts. At *NRG1*, significant risks at rs16879552 and rs16879576 were only observed in HDRC (3%/4%/8% and 2%/3%/7% allele frequency in controls/AC-HSCR/HDRC with ORs of 1.3 (95% CI: 0.85-1.88) vs. 3.1 (95% CI: 2.18-4.48) and 1.7 (95% CI: 1.09-2.51) vs. 3.6 (95% CI: 2.44-5.36) in AC-HSCR and HDRC for rs16879552 and rs16879576, respectively), in accord with results from our previous study of overlapping AC-HSCR cases^11^ (Table S6). At the *RET* locus, there were several variants with differential effects on HSCR risk between AC-HSCR and HDRC, including five with significant risk in AC- HSCR cases only (rs3758514, rs788243, rs7069590, rs12764797 and rs2472737), five with significant risk in HDRC cases only (rs3004255, rs4949062, rs3026693, rs1800860 and rs10793423), and at least five with substantially increased risk in HDRC as compared to AC- HSCR (rs2435357, rs2506024, rs2505515, rs2505513 and rs7074964) (Table S8). Overall, these results demonstrate that generally we detect a greater magnitude of associations in HDRC as compared to AC-HSCR, i.e., in samples enriched for simplex S-HSCR cases. For further analyses, any variant with statistically significant association in AC-HSCR, HDRC or AC-HSCR and HDRC combined cases was considered to be associated with HSCR risk.

### Risk of HSCR is additive across variants and genes

Having observed multiple single variant associations at *RET* (28 variants; Tables 2, S8; Figure S7), *SEMA3* (8 variants; Tables 2, S8; Figure S8) and *NRG1* (2 variants; Tables 2, S8), we next assessed their cumulative effects, within and across loci, on HSCR risk (Tables 3, S9-S10; Figure 1, S9). Here, we analyzed the two collections as one sample. For within locus analysis, given the rarity of risk alleles at the significant *NRG1* variants rs16879552 and rs16879576, we didn’t assess their effect. Risk alleles across all significant variants within *RET* and *SEMA3* loci were counted in each individual with neighboring classes combined to generate five non-overlapping bins to reduce numbers of tests and to have similar sample sizes across bins in controls (Tables S9, S10). Similarly, for analysis across loci, risk alleles at all significant variants were counted in each individual and combined to generate five non-overlapping bins (Table 3).

**Figure 1:**
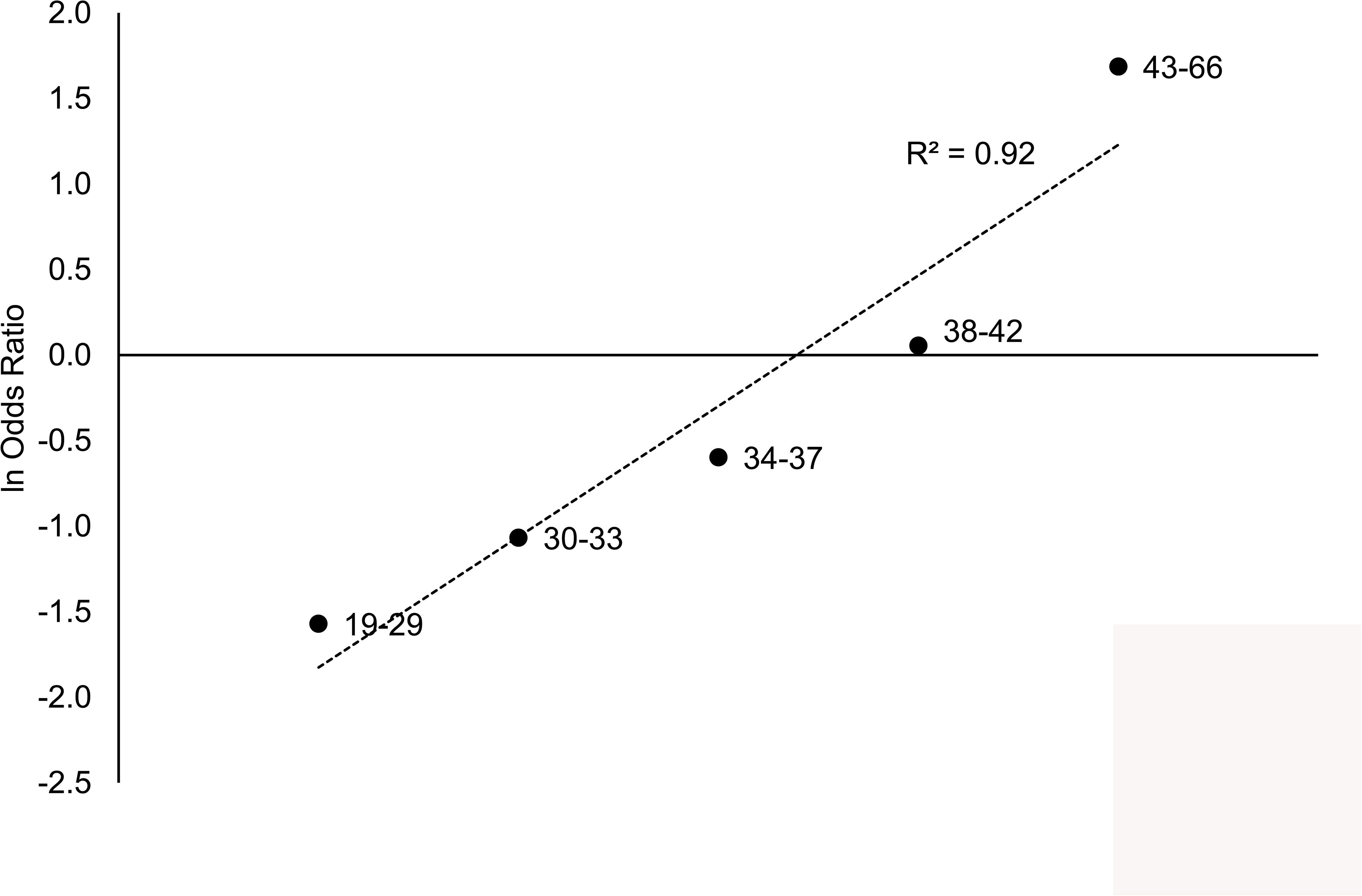
Logistic function relating HSCR risk with the total number of risk alleles at *RET*, *SEMA3* and *NRG1* loci. The logarithm of odds ratios (Y-axis) for binned risk allele counts (X- axis) based on significant variants across the three loci are shown along with the best linear fit (dashed line).

At *RET*, the total risk allele dosage was significantly associated with HSCR risk (Table S9; *χ*^2^=134.7, *P*=3.9×10^-28^; Figure S9) with the first three bins showing significant protection (OR=0.3/0.4/0.5 and *P*=3.2×10^-8^/2.8×10^-6^/6.7×10^-5^, for 17-23, 24-26 and 27-29 risk alleles, respectively) and the highest risk alleles bin having significantly increased disease risk (OR=4.9, *P*=2.8×10^-26^ for 35-52 risk alleles). The logistic function describing the relationship between HSCR risk and *RET* risk allele counts explains 80% of risk variation (Figure S9). We estimated the population penetrance (frequency of being affected given a genotype), which ranged from 4.9 cases to 40.0 cases for the lowest (17-23 risk alleles) to the highest (35-52 risk alleles) bins, respectively (Table S9). These values translate to a population incidence difference ranging from ∼1/20,000 to ∼1/2,500 live births. At *SEMA3*, although there was a trend for an analogous association with risk allele dosage (Table S10; *χ*^2^=11.9, *P*=0.018; Figure S9), only two risk alleles bins (1-3 risk alleles, OR=0.7, *P*=0.014; 8-9 risk alleles, OR=1.5, *P*=0.019) showed a trend towards statistical significance. The logistic function describing the relationship between HSCR risk and *SEMA3* risk allele counts explains 78% of risk variation (Figure S9). Counting risk alleles across the three loci led to an expected association between total risk allele dosage and HSCR risk (Table 3; *χ*^2^=156.4, *P*=8.5×10^-33^; Figure 1), which, however, was more significant when compared to risk from the *RET* locus alone (Table S9; *χ*^2^=134.7, *P*=3.9×10^-28^; Figure S9). The risk for HSCR from these variants increased from 0.2 to 5.4, a 26-fold change, as the bin size increased from 19-29 to 43-66 risk alleles. Also, the logistic function describing the relationship between HSCR risk and risk allele counts across the three loci explains 92% of risk variation, more than explained from *RET* alone (Figures 1, S9). The estimated population penetrance ranged from 3.7 cases to 45.4 cases for the lowest (19-29 risk alleles) to the highest (43-66 risk alleles) bins, respectively (Table 3). These values translate to a population incidence difference ranging from ∼1/27,000 to ∼1/2,200 live births. Collectively, these results show a clear additive effect of *RET* risk variants on HSCR risk and provide evidence for minor additional effects from *SEMA3* and *NRG1* risk variants.

Next, we wanted to assess how the combined (polygenic) risk from these variants varied across HSCR subjects. Polygenic risk score (PRS) for each HSCR subject was calculated and varied from 5.12 to 16.04 (median=10.66) and from 6.47 to 16.46 (median=11.28) in AC-HSCR and HDRC cohorts, respectively. Although, the PRS variation followed normal distributions in both cohorts with mean values of 10.60 and 11.19 for AC-HSCR and HDRC, respectively, there was indeed a small yet significant difference in the two sample means (z=2.63; *P*=0.009; Figure 2). We expect the difference in composition of HSCR subtypes between the two cohorts (Table 1) to be a major factor leading to demonstrable variation in polygenic risk.

**Figure 2:**
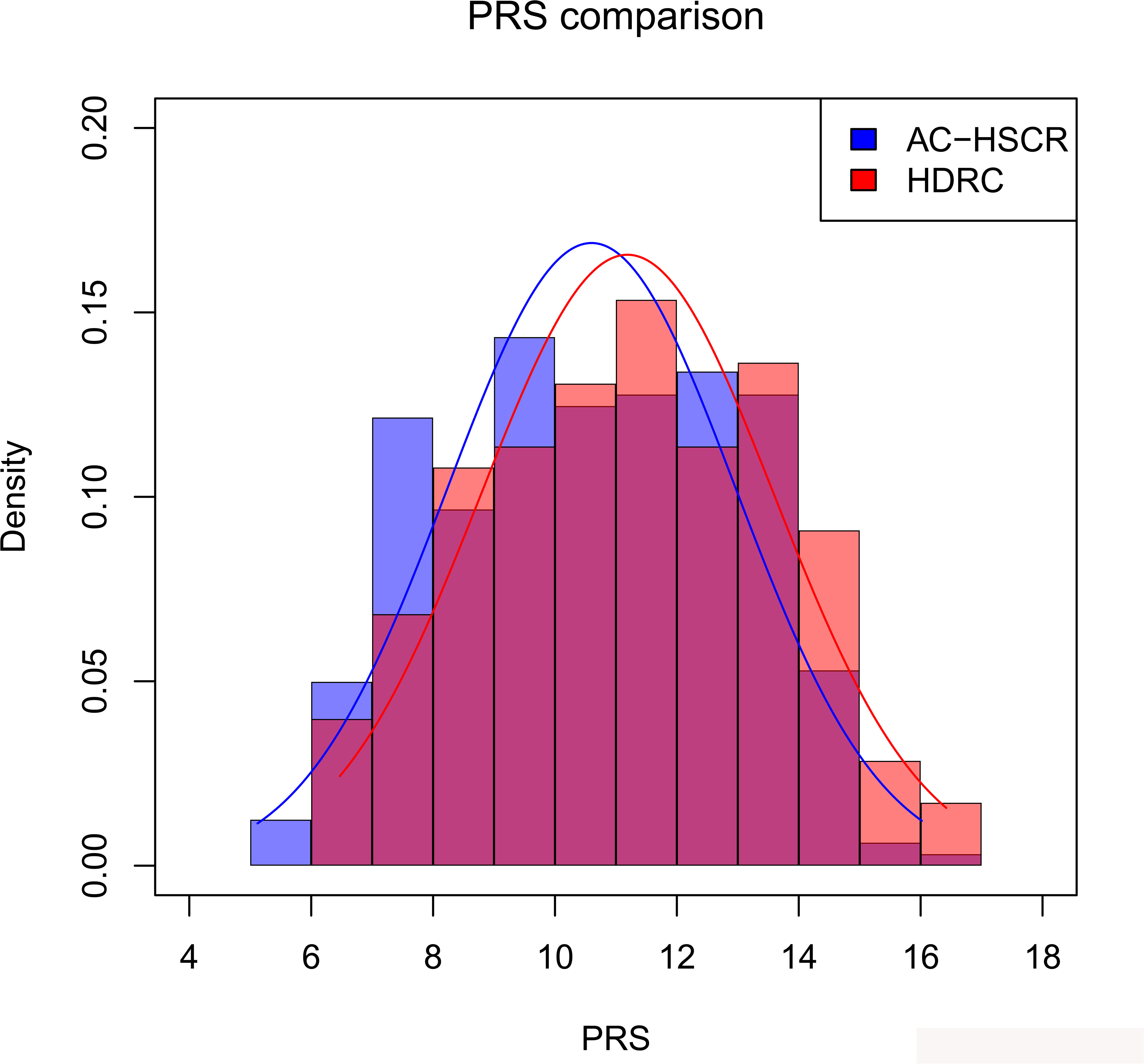
Polygenic risk score distributions. Empirical frequency distributions of polygenic risk scores (PRS) in AC-HSCR (blue) and HDRC (red) HSCR subjects are plotted along with their corresponding fitted normal distributions.

## Discussion

Evidence in the literature,^6–9,11,26,27^, and the far more extensive data reported here (Table 2, Table S8), more than adequately confirms the role of multiple CRE polymorphisms at *RET*, *SEMA3* and *NRG1* on HSCR susceptibility in European ancestry subjects. The most parsimonious hypothesis is that these gut enhancer variants reduce gene expression at *RET*, *SEMA3* and *NRG1* to increase the risk of aganglionosis, with risk increasing as a logistic function of the numbers of risk alleles (Figure 1). Although the reduction of *RET* expression through its polymorphic enhancer risk variants has been demonstrated by us,^12^ this remains to be to be shown for *SEMA3* and *NRG1*. A first question is whether marked expression reduction at *RET* leads to HSCR given its large impact relative to *SEMA3* and *NRG1*? This is unlikely because disease also occurs in non-risk allele homozygotes at *RET*. We have recently demonstrated,^10^ following years of indirect evidence from segregation,^2^ linkage^18,19^ and association studies,^11^ that individual patients harbor multiple rare risk coding variants across different genes. Here, we show the same feature but across common CRE risk variants. This prompts the second question: how does an enteric neuron ‘count’ risk variants or recognize expression change across genes to display its appropriate phenotype?

The mechanical and technical aspects of mapping variants to disease does not, however, clarify the underlying ‘counting’ mechanism even when we know that disease results from gene expression alteration. For understanding disease pathophysiology, we need to determine how much expression alteration is required before aganglionosis results. Clearly, single variants, the units of mapping, show disease association. But this does not imply that the variant alone leads to disease, indeed, it likely doesn’t because its genetic effect is usually small. Further, even 50% gene expression reduction at *RET* doesn’t always lead to a clinical phenotype.^8^ Thus, there is an expression threshold at each gene before disease results and this threshold is unlikely to be reached by single variants or perhaps even single genes or their enhancers. Consequently, multiple functional variants are required to act together to have a sufficient effect on gene expression to reach this threshold; the more extreme the threshold the larger the genetic effect whether from one or many variants.

The genetic consequences of this hypothesis implies that these risk variants are ‘counted’ together within a locus. One possibility is that they physically exist on a single chromosome in *cis* when they exert an effect, irrespective of whether these variants are in LD or not. Indeed, the causal genetic variants may be correlated, leading to the suspicion that they do not impart independent effects, when it is precisely this dependence (accumulation) that leads to a ‘jackpot’ effect on gene expression by that haplotype. Note that LD is not required but that its existence between variants does not invalidate their individualistic role on gene expression.

How then are the effects counted across genes? Current data suggests that the *RET*, *SEMA3* and *NRG1* effects on HSCR are additive, suggesting that their expression effects are independent; but they need not be. As we have shown for *RET*,^12^ developmental genes function within a GRN and the effects of a set of variants on one gene affects the expression of other network genes. Thus, combinations of alleles at these three genes may impart greater risk through affecting all genes in the network. Proving this hypothesis using human genetic data alone is going to be arduous because HSCR is a rare disorder and difficult to collect thousands of samples. In contrast, mouse models with enhancer variants at *Ret*, *Sema3* and *Nrg1* can be used to test the contention that even with variants that reduce gene expression at developmentally important disease genes, disease only results from specific combinations of genotypes across these loci depending on the architecture of the GRN and three-dimensional chromatin contacts. Such a hypothesis may be sufficient to explain the need for multiple genes in a disease, multiple variants in these genes, small genetic effects of disease-associated alleles, the familiality of the disorder and its reduced penetrance.

## Supplementary Methods

### Patient samples

Diagnosis of HSCR was based on surgical reports, pathological examination of rectal biopsies or other medical records. Patients were categorized by segment length of aganglionosis into short-segment HSCR (S-HSCR; aganglionosis limited up to the upper sigmoid colon), long-segment HSCR (L-HSCR; aganglionosis up to splenic flexure and beyond) or total colonic aganglionosis (TCA; aganglionosis of the entire large intestine); the segment length was indeterminable from records in ∼30% of cases. We used genomic DNA isolated from peripheral blood, buccal swabs or saliva samples from participants using standard protocols.

Genotyping was attempted in a total of 2,205 unique DNA samples from 737 families.

### Variant selection

For the *RET* and *SEMA3* loci, target regions for variant selection were identified based on our published HSCR genome-wide association study (GWAS) signals^1^ and flanking recombination hotspots data from HapMap^2^ (ftp://ftp.hapmap.org/hapmap/recombination/2008-03_rel22_B36/rates/), as implemented in LocusZoom (http://locuszoom.org/). This led to a target region of ∼575 kb at *RET* (chr10:43,222,994 – 43,797,994; hg19) (Figure S1) and ∼756 kb at *SEMA3* (chr7:83,953,750 – 84,709,831; hg19) (Figure S2). We used the 1000 Genomes^3^ non-Finnish European (NFE) ancestry (*n*=404) genotype data to select 1,227 and 1,056 common variants (minor allele frequency (MAF) >10%) within the *RET* and *SEMA3* loci, respectively. Next, at each locus we retained only variants that overlapped putative enhancer marks from public epigenomic databases,^4,5^ as described earlier:^6^ fetal large intestine DNaseI hypersensitivity (DHS), H3K4me1 and H3K27ac peaks, SK-N-SH retinoic acid treated DHS peaks,^4,5^ and untreated ATAC-peaks (unpublished data). This retained 167 variants at the *RET* locus and 67 at the *SEMA3* locus. In our prior HSCR GWAS^1^ 38 variants reached genome-wide significance at the *RET* locus, of which 10 demonstrated allelic differences in enhancer activities in *in vitro* reporter assays (Chatterjee *et al.* manuscript). Of these 10, 7 were part of the 167 *RET* variants we filtered; we added the remaining 3 for a total of 170 variants at the *RET* locus (Dataset S1).

### Multiplex genotyping assay

We used the MassARRAY Assay Design Suite 2.0 software (https://agenacx.com/) to design custom multiplexed genotyping assays using iPLEX chemistry, following the manufacturer’s recommendations (Agena Bioscience, San Diego, CA, USA). All steps of the genotyping assay (adjustment of extension primers, PCR amplification, shrimp alkaline phosphatase treatment, iPLEX extension reaction, desalting of iPLEX extension reaction product, dispensing of samples to a SpectroCHIP array, data acquisition on the MassARRAY analyzer and data processing by Typer software) were performed following the manufacturer’s recommendations (Agena Bioscience). In the first round of assay design, two multiplexed assay pools, #1 with 32 variants and #2 with 30 variants, were designed and evaluated using reference DNA samples. Four variants in each pool failed in genotyping (due to amplification or extension failures) and were replaced with a best proxy variant from the filtered sets of variants when available; if not, we used the best replacement from the larger sets of common variants (based on LD) to generate two new multiplexed assay pools, #1 with 33 variants and #2 with 28 variants (Dataset S1). Study samples were genotyped using these re-designed multiplexed assay pools.

Dispensing of desalted iPLEX reaction products to SpectroCHIP arrays, data acquisition on the MassARRAY analyzer and data processing by Typer software were performed at ReproCELL USA Inc. (Beltsville, MD, USA).

### Variant genotyping quality control (QC)

Genotyping of 2,205 unique DNA samples was carried out in six 384-well plates/chips for each multiplexed pool. Before variant quality control (QC), 3 and 2 samples in pools #1 and #2, respectively, were removed due to dispensing failure. Then, 2 variants in pool #1 (rs7458400 and rs7900769) and 1 variant in pool #2 (rs3097563) were dropped due to high genotyping missing rates (>10%) (Dataset S1, Table S1). Next, 2 variants, both in pool #2 were removed: one was nearly monomorphic (rs3026696, with a MAF of ∼15% in 1000 Genomes NFE ancestry subjects) and the other (rs4948695) had almost no heterozygous calls, both indicative of poor extension, mass separation and clustering (Dataset S1). We also removed samples with >10% missing genotypes within each pool (Table S2). Due to failures at various steps, from assay design to variant QC, 14 out of 170 filtered variants at *RET* and 1 out of 67 filtered variants at *SEMA3* were not studied (Dataset S1).

### Statistical genetic analyses

We used publicly available allele count data from the Genome Aggregation Database (gnomAD)^7^ (http://gnomad-old.broadinstitute.org/), based on whole-genome sequencing of NFE ancestry subjects. This is a large control population dataset with allele counts ranging from 14,172 to 15,008 across the 56 tag variants (median allele count=14,953). Case-control allelic association analysis was performed using standard contingency *χ^2^* tests; standard methods were used for calculations of OR, their confidence limits and statistical significance of association (OR=1). Pairwise LD between all tag variants at each locus was estimated in HSCR cases and 1000 Genomes^3^ NFE ancestry subjects using Haploview^8^. For analyzing the combined effects of multiple variants at each locus, we only considered variants significantly associated with HSCR risk and counted the total number of risk alleles within a locus in each individual (case or 1000 Genomes NFE ancestry control). Risk allele counts were binned into non-overlapping bins keeping similar proportions of controls in each bin.

## Acknowledgments

We wish to thank the numerous patients, their families, referring physicians, nurses and genetic counselors who have contributed to these studies, and Erick Kaufmann, Jennifer (Scott) Bubb, Maura Kenton, Julie Albertus and Magan Trottier for family ascertainment, family studies and genetic counseling. The studies reported here were partially supported by a grant from the US National Institutes of Health (MERIT Award HD28088 to A.C.).

**Hirschsprung Disease Research Collaborative (HDRC)**

**Corresponding author**

Aravinda Chakravarti^1,2,#^

**Steering Committee**

Courtney Berrios^1,3,#^

Aravinda Chakravarti^1,2,#^

Philip Frykman^4^

Cheryl Gariepy^5^

Raj Kapur^6^

Jacob Langer^7^

**Sample Collection**

Jeffrey Avansino^8^ (Principal Investigator), Courtney Berrios^1,3,#^, Andrea Bischoff^9^ (Principal Investigator), Aravinda Chakravarti^1,2,#^ (Principal Investigator), Nicole Chandler^10,***^ (Principal Investigator), Robert Cina^11^ (Principal Investigator), Daniel DeUgarte^12^ (Principal Investigator), Megan Durham^13^ (Principal Investigator), Jason Frischer^14,***^ (Principal Investigator), Philip Frykman^4^ (Principal Investigator), Samir Gadepalli^15^,*** (Principal Investigator), Cheryl Gariepy^5^ (Principal Investigator), Ankush Gosain^16^ (Principal Investigator), Gunadi^17^ (Principal Investigator), Raj Kapur^6^ (Principal Investigator), Jacob Langer^7^ (Principal Investigator), Monica Lopez^18^ (Principal Investigator), Lisa McMahon^19^ (Principal Investigator), Suyin Lum Min^20,***^ (Principal Investigator), Hector Monforte^10,***^, Isam Nasr^21^, Dorothy Rocourt^22^ (Principal Investigator), David Rodeberg^23^ (Principal Investigator), Michael Rollins^24^ (Principal Investigator), Robert Russell^25^ (Principal Investigator), Payam Saadai^26^ (Principal Investigator), Lois Sayrs^19^, Donald Shaul^27,***^ (Principal Investigator), Douglas Tamura^28^ (Principal Investigator), Richard Weiss^29^ (Principal Investigator), Jia Yan^1,2,#^

^1^McKusick-Nathans Institute of Genetic Medicine, Johns Hopkins University School of Medicine, Baltimore, MD 21205, USA; ^2^Center for Human Genetics and Genomics, New York University School of Medicine, New York, NY 10016, USA; ^3^Childrens’s Mercy Research Institute, Kansas City, MO 64108, USA; ^4^Department of Pediatric Surgery, Cedars-Sinai Medical Center, Los Angeles, CA 90048, USA; ^5^Department of Gastroenterology, Hepatology & Nutrition, Nationwide Children’s Hospital, Columbus, OH 43205, USA; ^6^Department of Laboratories Pathology, Seattle Children’s Hospital, Seattle, WA 98105, USA; ^7^Department of General and Thoracic Surgery, The Hospital for Sick Children, Toronto, ON M5G 1X8, Canada; ^8^Department of Pediatric Surgery, Seattle Children’s Hospital, Seattle, WA 98105, USA; ^9^Children’s Hospital Colorado, University of Colorado School of Medicine, Aurora, CO 80045, USA; ^10^All Children’s Hospital, St. Petersburg, FL 33701, USA; ^11^Department of Surgery, Medical University of South Carolina College of Medicine, Charleston, SC 29425, USA; ^12^Department of Surgery, University of California Los Angeles School of Medicine, Los Angeles, CA 90095, USA; ^13^Children’s Healthcare of Atlanta, Emory University School of Medicine, Atlanta, GA 30322, USA; ^14^Cincinnati Children’s Hospital Medical Center, University of Cincinnati, Cincinnati, OH 45229, USA; ^15^Pediatirc Surgery, Michigan Medicine, Ann Arbor, MI 48109, USA; ^16^Le Bonheur Children’s Hospital, University of Tennessee Health Science Center, Memphis, TN 38103, USA; ^17^Department of Surgery, University of Gadjah Mada, Yogyakarta 55281, Indonesia; ^18^Pediatric Surgery, Texas Children’s Hospital, Houston, TX 77030, USA; ^19^Department of Surgery, Phoenix Children’s Hospital, Phoenix, AZ 85016, USA; ^20^The Children’s Hospital of Winnipeg, University of Manitoba, Winnipeg, MB R3A 1S1, Canada; ^21^Department of Surgery, Johns Hopkins University School of Medicine, Baltimore, MD 21205, USA; ^22^Pediatric Surgery, Penn State Health Children’s Hospital, Hershey, PA 17033, USA; ^23^Department of Surgery, East Carolina University School of Medicine, Greenville, NC 27834, USA; ^24^Department of Surgery, University of Utah School of Medicine, Salt Lake City, UT 84132, USA; ^25^Department of Surgery, University of Alabama at Birmingham School of Medicine, Birmingham, AL 35233, USA; ^26^Department of Surgery, University of California, Davis School of Medicine, Sacramento, CA 95817, USA; ^27^Kaiser Permanente Southern California, Los Angeles, CA 90027, USA; ^28^Valley Children’s Hospital, Madera, CA 93636, USA; ^29^Pediatric Surgery, Connecticut Children’s Medical Center, Hartford, CT 06106, USA.

***Site no longer part of the HDRC

^#^Author of the manuscript; everyone else is a contributor

## Conflicts of Interest

The authors have no conflicts of interest to report.

---

**Figure S1:**
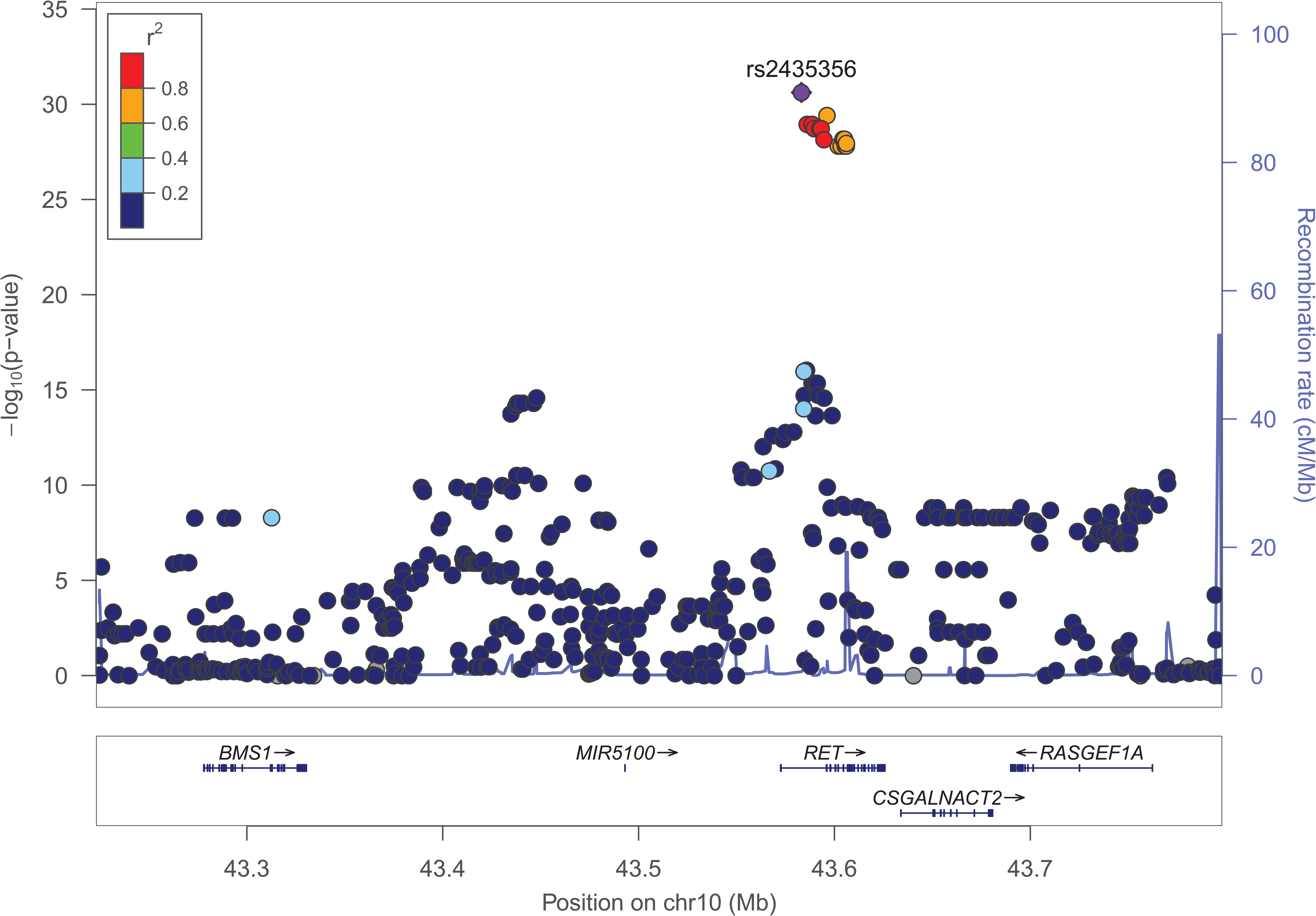
Regional association plot of variants at the *RET* locus in HSCR GWAS.^1^. The X-axis is the genomic interval annotated with gene models, the left Y-axis the statistical significance of association (-log_10_p), and the right Y-axis the recombination rate (blue trace, cM (centiMorgan)/Mb(megabase)) based on HapMap samples.^2^ The most significant SNP, rs2435356, is shown as a purple circle. All other variants are color-coded based on their LD (*r^2^*) with the index variant. The plot was generated with *LocusZoom* (http://locuszoom.org/).

**Figure S2:**
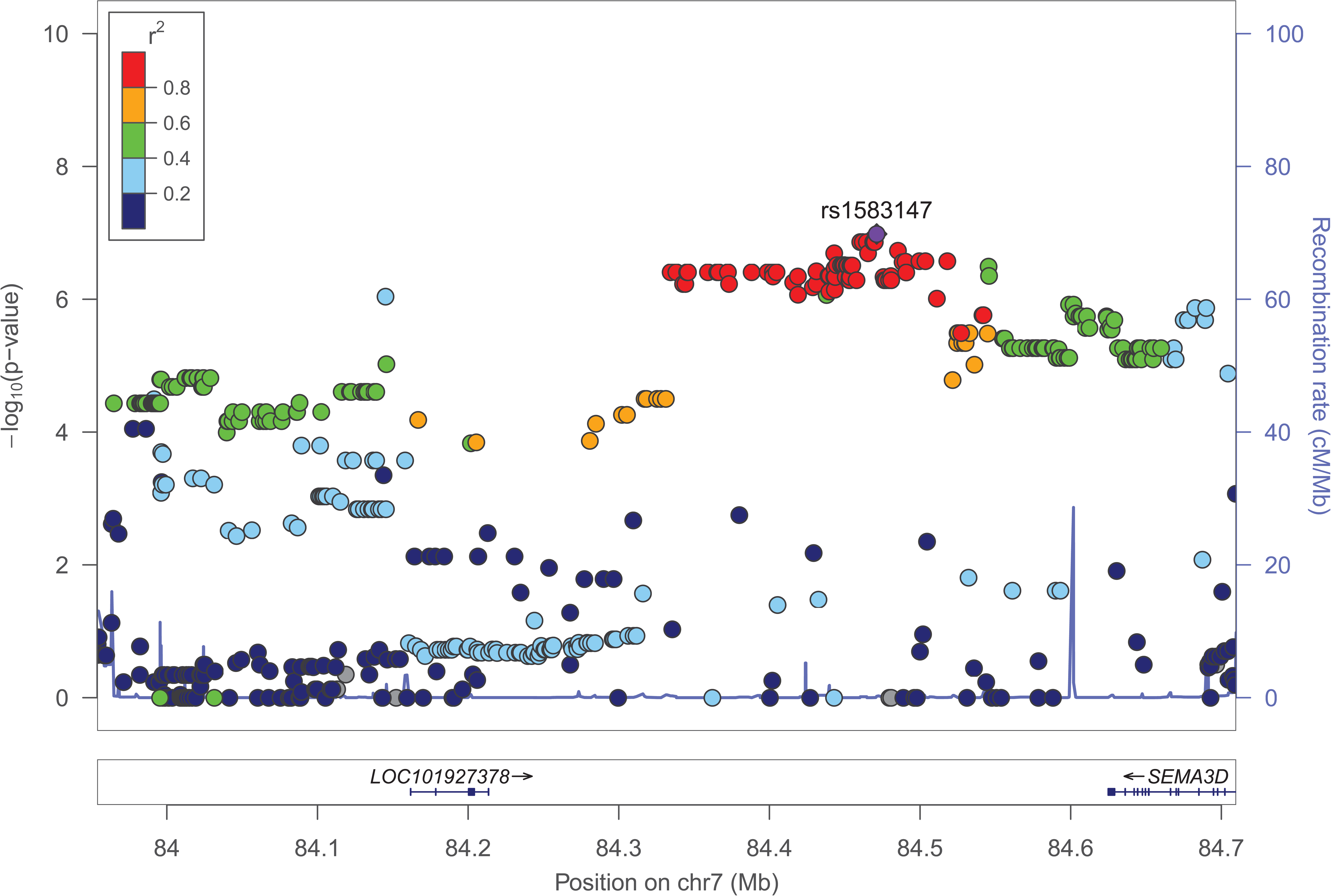
Regional association plot of variants at the *SEMA3* locus in HSCR GWAS.^1^. The X-axis is the genomic interval annotated with gene models, the left Y-axis the statistical significance of association (-log_10_p), and the right Y-axis the recombination rate (blue trace, cM (centiMorgan)/Mb(megabase)) based on HapMap samples.^2^ The most significant SNP, rs1583147, is shown as a purple circle. All other variants are color-coded based on their LD (*r^2^*) with the index variant. The plot was generated with *LocusZoom* (http://locuszoom.org/).

**Figure S3:**
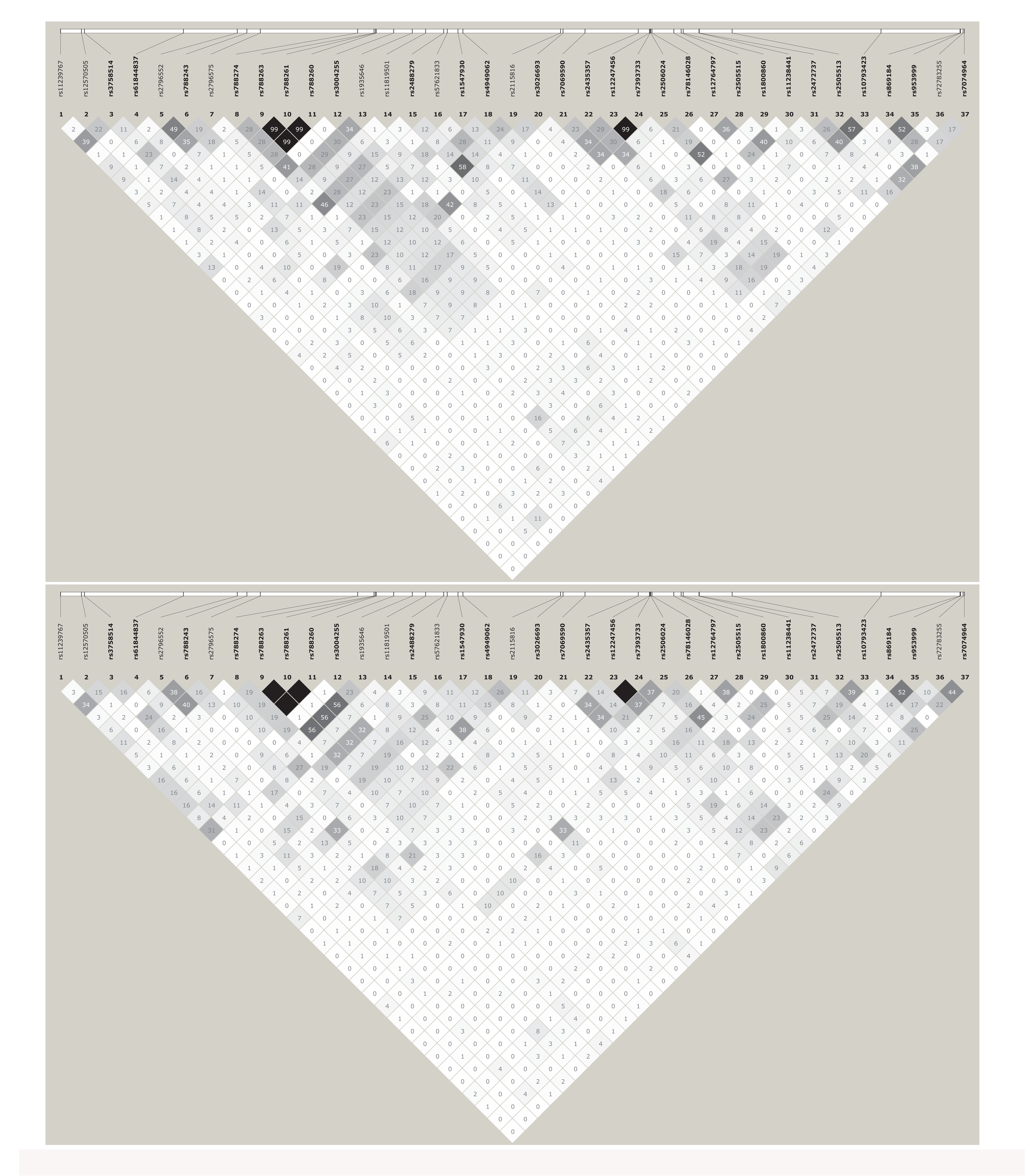
LD matrix for tag variants at the *RET* locus in HSCR cases and controls. All pairwise LD (*r^2^*) among HSCR cases (n=583; *top*) and 1000 Genomes^3^ NFE ancestry controls (n=404; *bottom*) for all tag variants at the *RET* locus was calculated and plotted using *Haploview*^8^ (https://www.broadinstitute.org/haploview/downloads). Variants with significant association to HSCR risk in case-control analysis are shown in bold.

**Figure S4:**
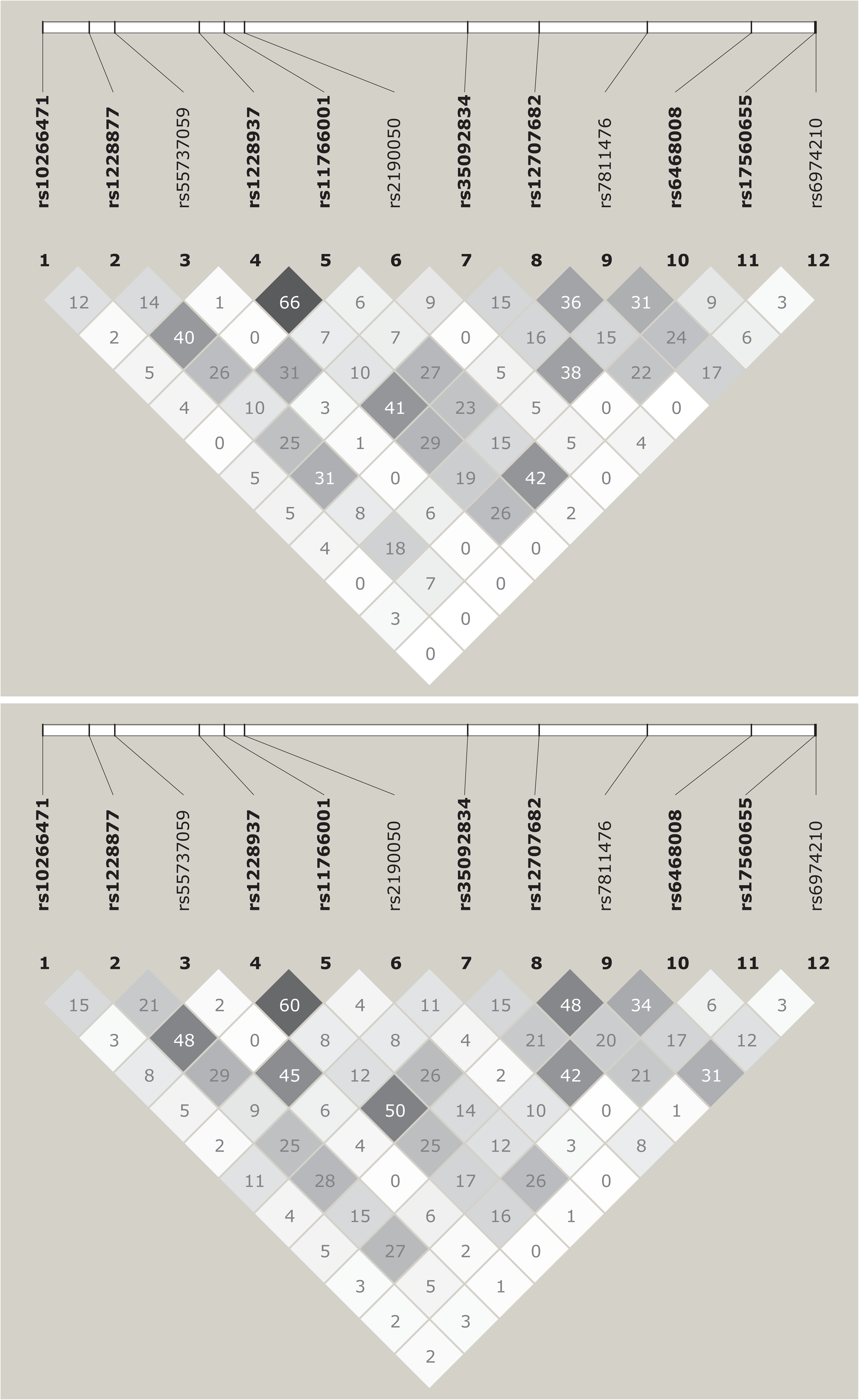
LD matrix for tag variants at the *SEMA3* locus in HSCR cases and controls. All pairwise LD (*r^2^*) among HSCR cases (n=583; *top*) and 1000 Genomes^3^ NFE ancestry controls (n=404; *bottom*) for all tag variants at the *SEMA3* locus was calculated and plotted using *Haploview*^8^ (https://www.broadinstitute.org/haploview/downloads). Variants with significant association to HSCR risk in case-control analysis are shown in bold.

**Figure S5:**
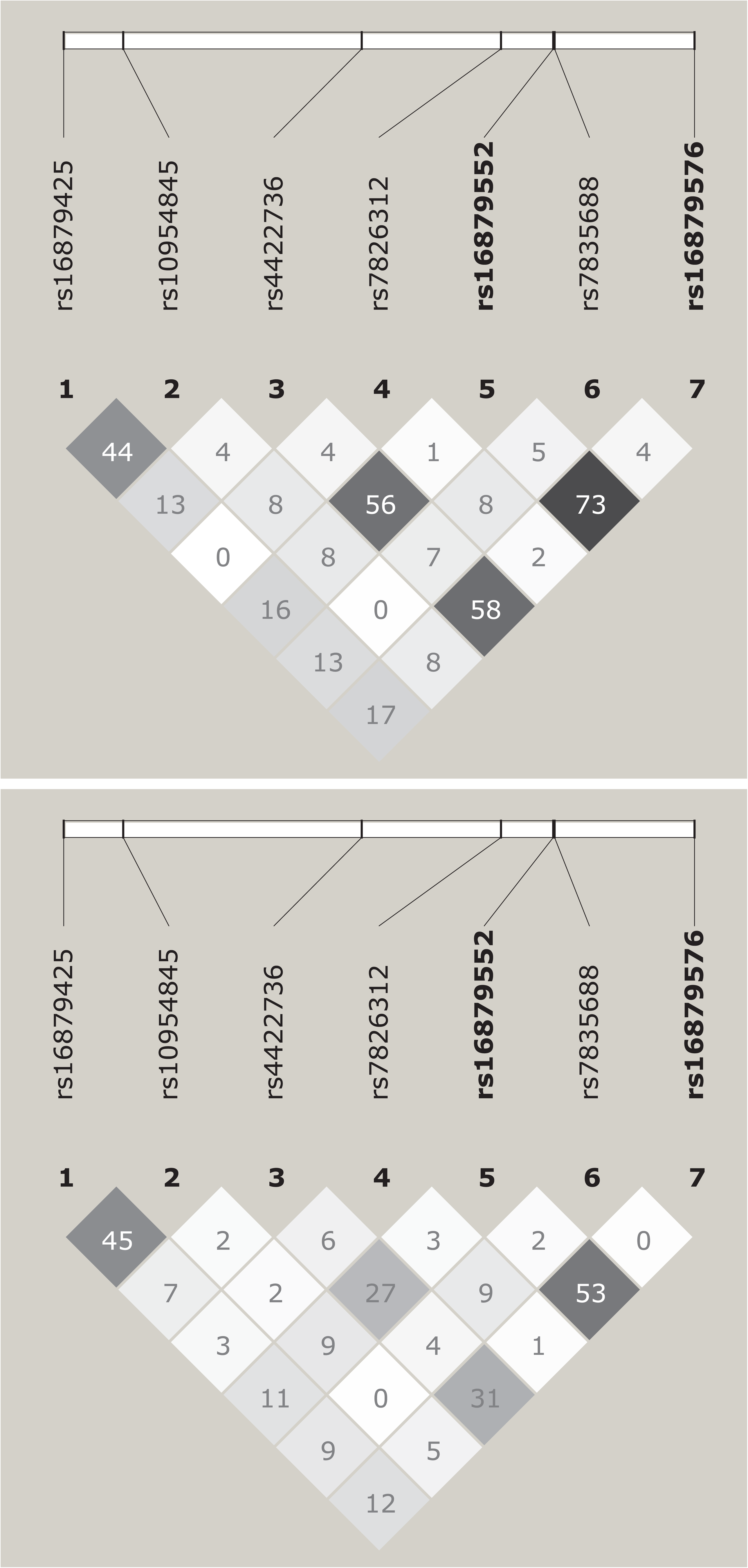
LD matrix for tag variants at the *NRG1* locus in HSCR cases and controls. All pairwise LD (*r^2^*) among HSCR cases (n=583; *top*) and 1000 Genomes^3^ NFE ancestry controls (n=404; *bottom*) for all tag variants at the *NRG1* locus was calculated and plotted using *Haploview*^8^ (https://www.broadinstitute.org/haploview/downloads). Variants with significant association to HSCR risk in case-control analysis are shown in bold.

**Figure S6:**
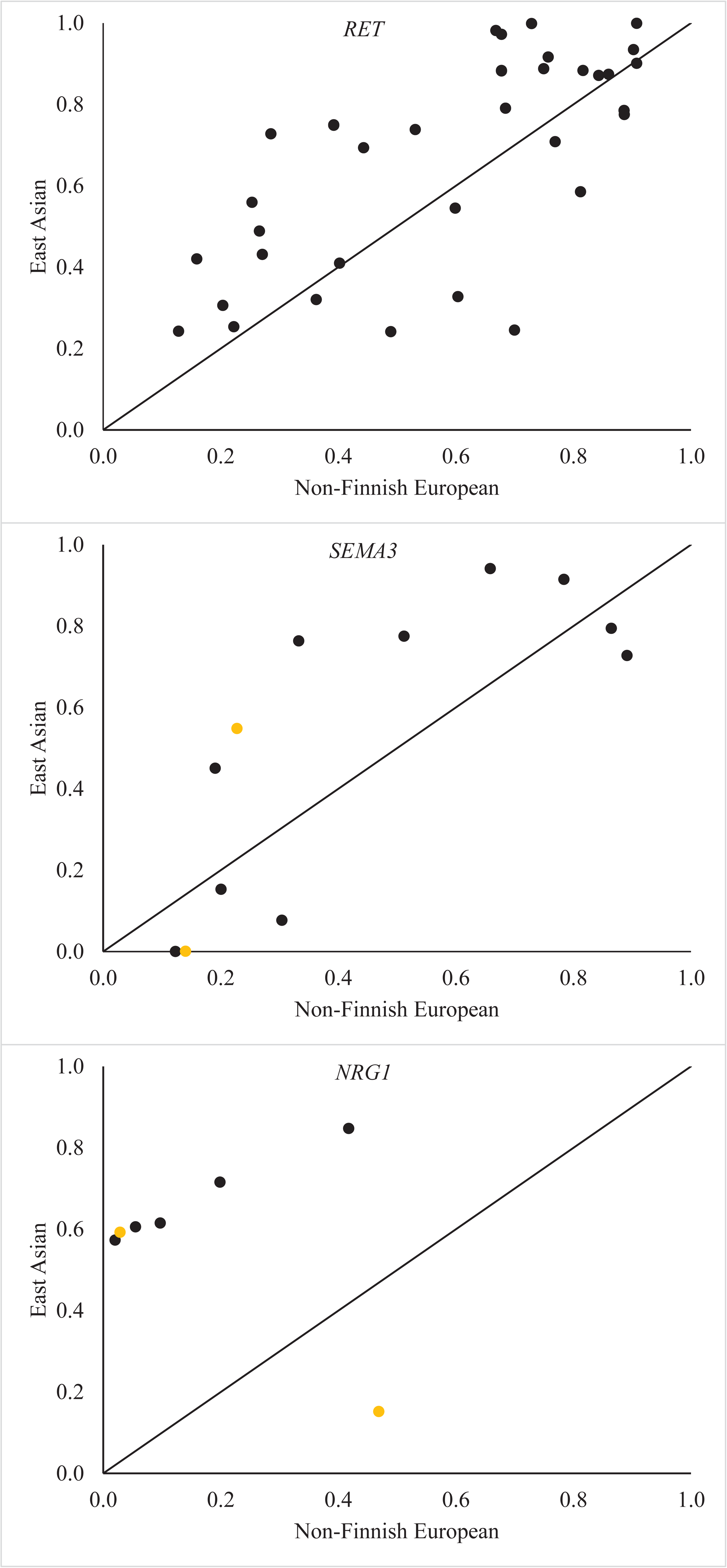
Allele frequency comparisons for tag variants between Non-Finnish European and East Asian ancestry gnomAD controls. X-Y scatter plots to compare allele frequencies of tag variants at *RET* (*n*=37; *top*), *SEMA3* (*n*=12; *middle*) and *NRG1* (*n*=7; *bottom*) between Non-Finnish European ancestry (X-axis) and East Asian ancestry (Y-axis) gnomAD controls.^7^ The sentinel hits at *SEMA3* in European ancestry HSCR GWAS^1^ and at *NRG1* in Asian ancestry HSCR GWAS^9^ are indicated by orange circles to highlight the contrast in observed allele frequencies.

**Figure S7:**
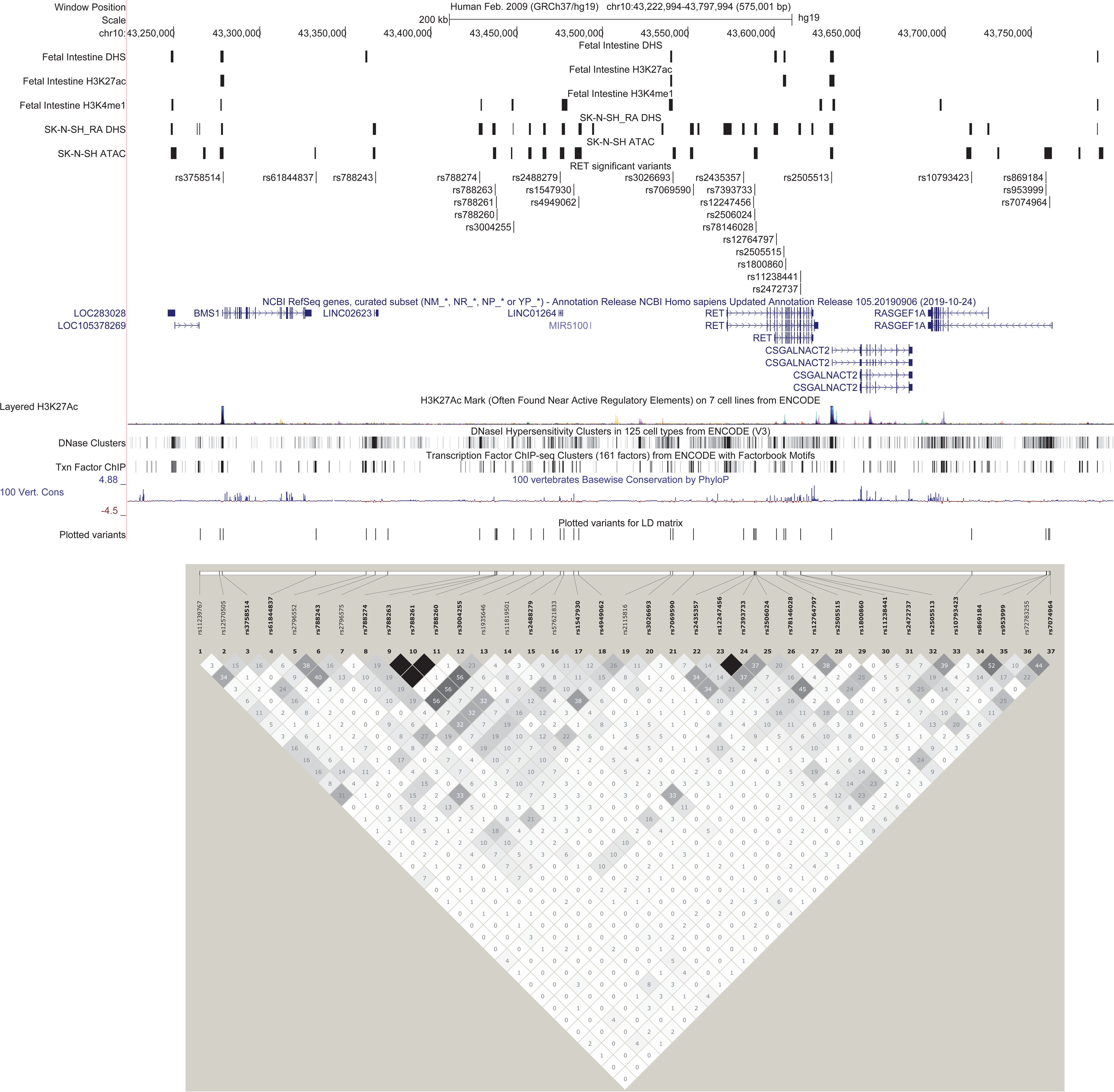
Genomic map of tag variants associated with overall HSCR risk at the *RET* locus. A 575 kb genomic segment annotated with tracks, showing (from top) human fetal intestine DNase-seq peaks (Fetal Intestine DHS (DNaseI hypersensitive sites))^5,6^, human fetal intestine H3K27ac ChIP-seq peaks (Fetal Intestine H3K27ac)^5,6^, human fetal intestine H3K4me1 ChIP-seq peaks (Fetal Intestine H3K4me1)^5,6^, DNase-seq peaks from SK-N-SH cells treated with retinoic acid (SK-N-SH_RA DHS)^4,6^, ATAC-seq peaks from SK-N-SH cells (SK-N-SH ATAC) (unpublished), tag variants with significant association to overall HSCR (RET significant variants), gene models (NCBI RefSeq genes), ENCODE integrated regulation H3K27ac marks (Layered H3K27Ac)^4^, ENCODE integrated regulation DNaseI hypersensitivity clusters (DNase Clusters)^4^, ENCODE integrated regulation Transcription factor ChIP-seq (Txn Factor ChIP)^4^, vertebrate basewise conservation by PhyloP (100 Vert. Cons)^10^ and the variants present in the LD matrix (Plotted variants). All pairwise LD (*r^2^*) among 1000 Genomes^3^ NFE ancestry controls (*n*=404) for all tag variants at the *RET* locus was calculated and plotted using *Haploview*^8^ (https://www.broadinstitute.org/haploview/downloads). Variants with significant association to HSCR risk in case-control analysis are shown in bold. The genomic map was generated using publically available and custom tracks in the UCSC genome browser (https://genome.ucsc.edu/).

**Figure S8:**
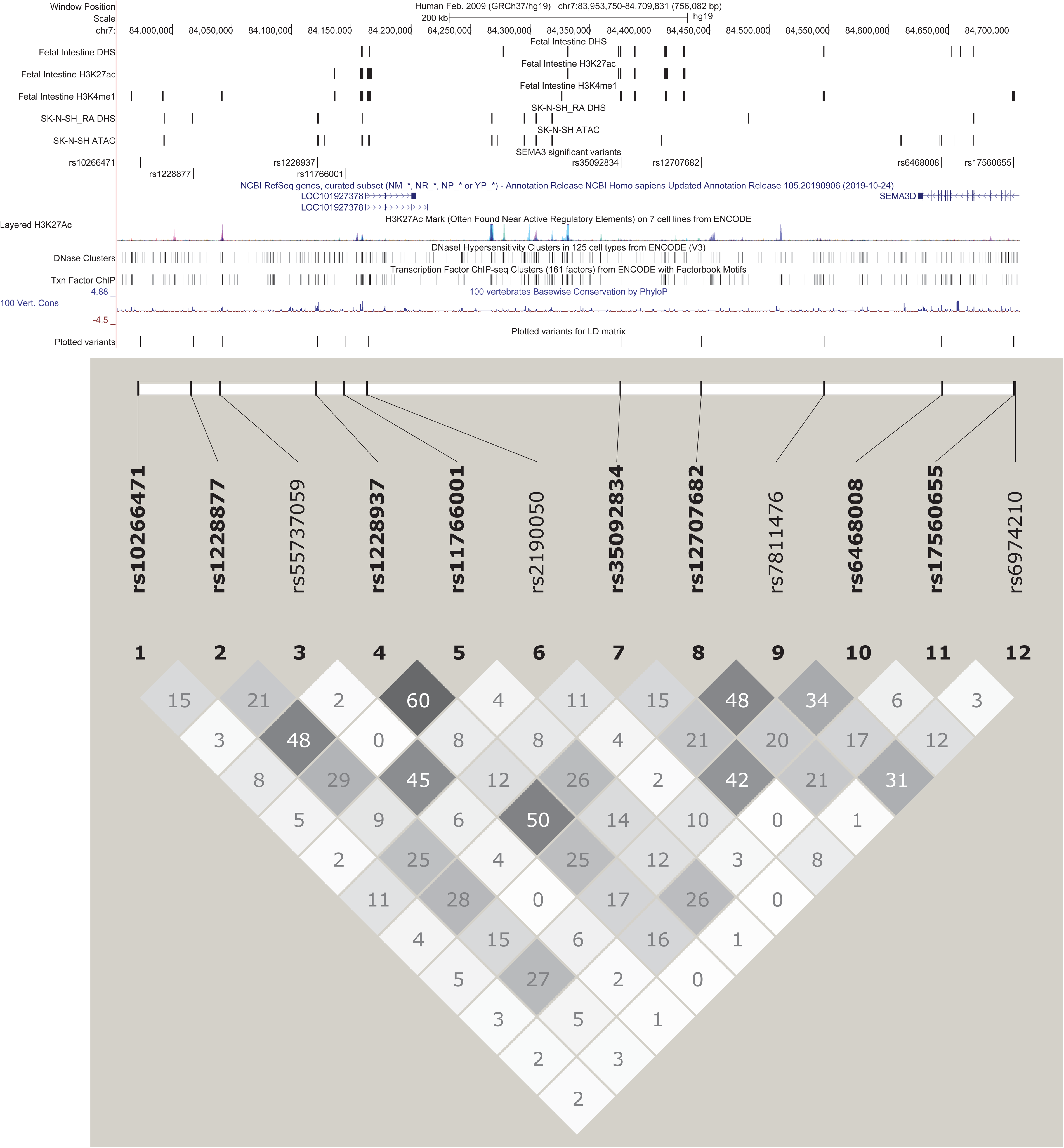
Genomic map of tag variants associated with overall HSCR risk at the *SEMA3* locus. A 756 kb genomic segment annotated with tracks, showing (from top) human fetal intestine DNase-seq peaks (Fetal Intestine DHS (DNaseI hypersensitive sites))^5.6^, human fetal intestine H3K27ac ChIP-seq peaks (Fetal Intestine H3K27ac)^5,6^, human fetal intestine H3K4me1 ChIP-seq peaks (Fetal Intestine H3K4me1)^5,6^, DNase-seq peaks from SK-N-SH cells treated with retinoic acid (SK-N- SH_RA DHS)^4,6^, ATAC-seq peaks from SK-N-SH cells (SK-N-SH ATAC) (unpublished), tag variants with significant association to overall HSCR (SEMA3 significant variants), gene models (NCBI RefSeq genes), ENCODE integrated regulation H3K27ac marks (Layered H3K27Ac)^4^, ENCODE integrated regulation DNaseI hypersensitivity clusters (DNase Clusters)^4^, ENCODE integrated regulation Transcription factor ChIP-seq (Txn Factor ChIP)^4^, vertebrate basewise conservation by PhyloP (100 Vert. Cons)^10^ and the variants present in the LD matrix (Plotted variants). All pairwise LD (*r^2^*) among 1000 Genomes^3^ NFE ancestry controls (*n*=404) for all tag variants at the *SEMA3* locus was calculated and plotted using *Haploview*^8^ (https:// www.broadinstitute.org/haploview/downloads). Variants with significant association to HSCR risk in case-control analysis are shown in bold. The genomic map was generated using publically available and custom tracks in the UCSC genome browser (https://genome.ucsc.edu/).

**Figure S9:**
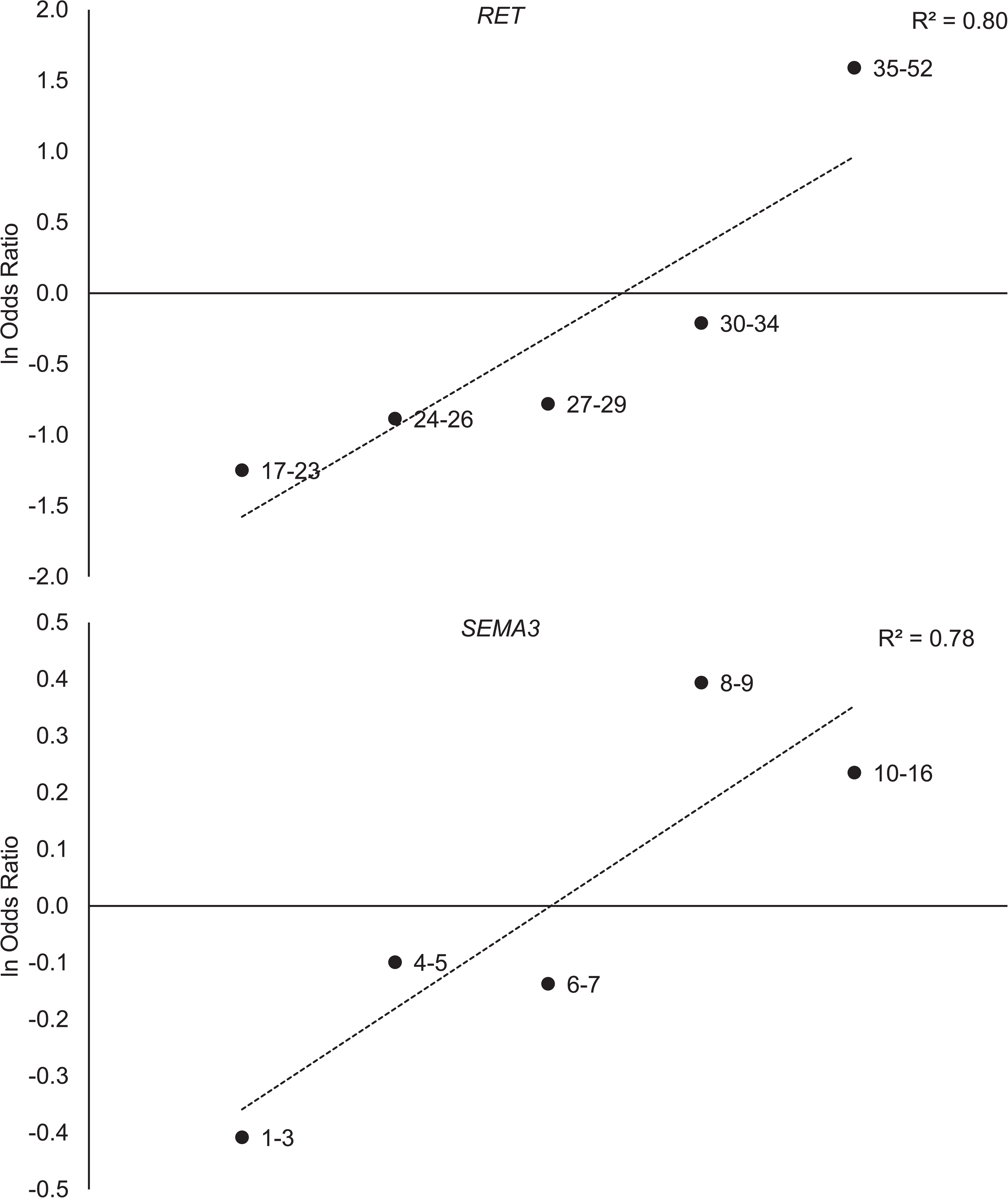
Logistic function relating HSCR risk with the total number of risk alleles at *RET* or *SEMA3* locus. The logarithm of odds ratios (Y-axis) for binned risk allele counts (X-axis) for *RET* (*top*) or *SEMA3* (*bottom*) significant variants are shown along with the best linear fit (dashed line).

**Table S1:** Percentage missing rate in each multiplexed genotyping pool by variant.

| **Pool #1 variant** | **Missing rate (%)** | **Pool #2 variant** | **Missing rate (%)** |
| --- | --- | --- | --- |
| rs10266471 | 3.1 | rs10793423 | 4.1 |
| rs10954845 | 3.6 | rs1228937 | 1.5 |
| rs11238441 | 3.5 | rs12570505 | 1.9 |
| rs11239767 | 3.5 | rs12707682 | 4.8 |
| rs11766001 | 7.0 | rs12764797 | 3.7 |
| rs11819501 | 3.4 | rs1547930 | 2.4 |
| rs12247456 | 4.6 | rs16879425 | 3.9 |
| rs1228877 | 2.7 | rs1800860 | 3.8 |
| rs16879552 | 3.5 | rs2115816 | 1.5 |
| rs16879576 | 3.4 | rs2190050 | 3.0 |
| rs17560655 | 2.5 | rs2435357 | 3.6 |
| rs1935646 | 3.0 | rs2488279 | 2.0 |
| rs2472737 | 3.8 | rs2505513 | 5.7 |
| rs2505515 | 3.7 | rs2796552 | 5.9 |
| rs2506024 | 5.7 | rs3026696 | 2.0 |
| rs2796575 | 2.2 | rs3097563 | 54.0 |
| rs3004255 | 4.5 | rs35092834 | 2.7 |
| rs3026693 | 2.9 | rs4422736 | 3.2 |
| rs3758514 | 3.1 | rs4948695 | 7.6 |
| rs4949062 | 4.0 | rs55737059 | 2.2 |
| rs6974210 | 3.9 | rs57621833 | 2.5 |
| rs7069590 | 2.7 | rs61844837 | 2.5 |
| rs7074964 | 5.8 | rs6468008 | 2.1 |
| rs72783255 | 3.2 | rs7393733 | 1.8 |
| rs7458400 | 31.2 | rs7811476 | 1.2 |
| rs78146028 | 3.5 | rs788261 | 2.0 |
| rs7826312 | 5.3 | rs788263 | 2.5 |
| rs7835688 | 4.0 | rs953999 | 1.6 |
| rs788243 | 4.5 |  |  |
| rs788260 | 4.6 |  |  |
| rs788274 | 4.1 |  |  |
| rs7900769 | 44.1 |  |  |
| rs869184 | 3.9 |  |  |

**Table S2:** Missing rate in each multiplexed genotyping pool across samples.

| **Pool #1 (31 variants)** |  | **Pool #2 (25 variants)** |  |
| --- | --- | --- | --- |
| **Number of**<br>**no calls** | **Number of**<br>**samples (%)** | **Number of**<br>**no calls** | **Number of**<br>**samples (%)** |
| 0 | 1,717 (78.0) | 0 | 1,832 (83.2) |
| 1 | 189 (8.6) | 1 | 157 (7.1) |
| 2 | 81 (3.7) | 2 | 70 (3.2) |
| 3 | 46 (2.1) | 3 or more | 144 (6.5) |
| 4 or more | 169 (7.7) |  |  |

**Table S3:**
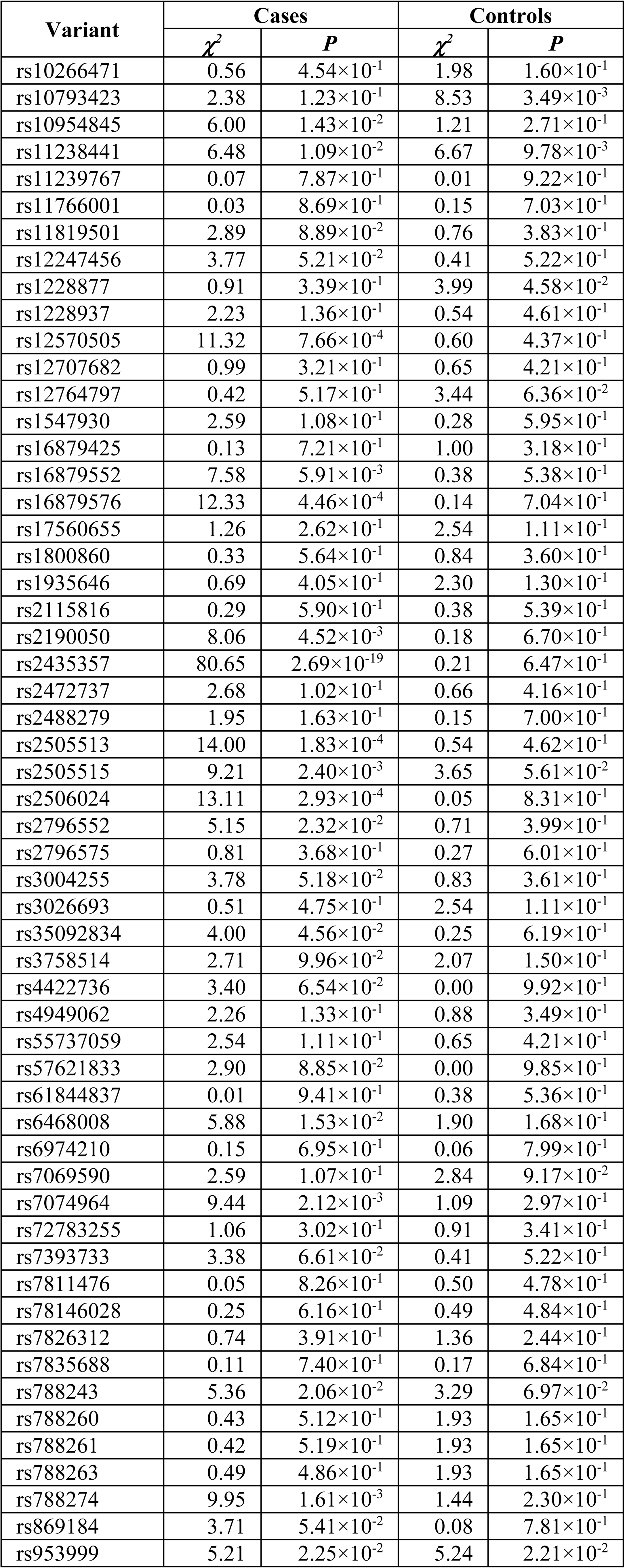
Tests of Hardy-Weinberg Equilibrium (HWE) by variant for HSCR cases and controls. Test statistic (*χ^2^*) and significance values (*P*) are provided.

| Variant | Cases |  | Controls |  |
| --- | --- | --- | --- | --- |
| | $\chi^2$ | $P$ | $\chi^2$ | $P$ |
| rs10266471 | 0.56 | $4.54 \times 10^{-1}$ | 1.98 | $1.60 \times 10^{-1}$ |
| rs10793423 | 2.38 | $1.23 \times 10^{-1}$ | 8.53 | $3.49 \times 10^{-3}$ |
| rs10954845 | 6.00 | $1.43 \times 10^{-2}$ | 1.21 | $2.71 \times 10^{-1}$ |
| rs11238441 | 6.48 | $1.09 \times 10^{-2}$ | 6.67 | $9.78 \times 10^{-3}$ |
| rs11239767 | 0.07 | $7.87 \times 10^{-1}$ | 0.01 | $9.22 \times 10^{-1}$ |
| rs11766001 | 0.03 | $8.69 \times 10^{-1}$ | 0.15 | $7.03 \times 10^{-1}$ |
| rs11819501 | 2.89 | $8.89 \times 10^{-2}$ | 0.76 | $3.83 \times 10^{-1}$ |
| rs12247456 | 3.77 | $5.21 \times 10^{-2}$ | 0.41 | $5.22 \times 10^{-1}$ |
| rs1228877 | 0.91 | $3.39 \times 10^{-1}$ | 3.99 | $4.58 \times 10^{-2}$ |
| rs1228937 | 2.23 | $1.36 \times 10^{-1}$ | 0.54 | $4.61 \times 10^{-1}$ |
| rs12570505 | 11.32 | $7.66 \times 10^{-4}$ | 0.60 | $4.37 \times 10^{-1}$ |
| rs12707682 | 0.99 | $3.21 \times 10^{-1}$ | 0.65 | $4.21 \times 10^{-1}$ |
| rs12764797 | 0.42 | $5.17 \times 10^{-1}$ | 3.44 | $6.36 \times 10^{-2}$ |
| rs1547930 | 2.59 | $1.08 \times 10^{-1}$ | 0.28 | $5.95 \times 10^{-1}$ |
| rs16879425 | 0.13 | $7.21 \times 10^{-1}$ | 1.00 | $3.18 \times 10^{-1}$ |
| rs16879552 | 7.58 | $5.91 \times 10^{-3}$ | 0.38 | $5.38 \times 10^{-1}$ |
| rs16879576 | 12.33 | $4.46 \times 10^{-4}$ | 0.14 | $7.04 \times 10^{-1}$ |
| rs17560655 | 1.26 | $2.62 \times 10^{-1}$ | 2.54 | $1.11 \times 10^{-1}$ |
| rs1800860 | 0.33 | $5.64 \times 10^{-1}$ | 0.84 | $3.60 \times 10^{-1}$ |
| rs1935646 | 0.69 | $4.05 \times 10^{-1}$ | 2.30 | $1.30 \times 10^{-1}$ |
| rs2115816 | 0.29 | $5.90 \times 10^{-1}$ | 0.38 | $5.39 \times 10^{-1}$ |
| rs2190050 | 8.06 | $4.52 \times 10^{-3}$ | 0.18 | $6.70 \times 10^{-1}$ |
| rs2435357 | 80.65 | $2.69 \times 10^{-19}$ | 0.21 | $6.47 \times 10^{-1}$ |
| rs2472737 | 2.68 | $1.02 \times 10^{-1}$ | 0.66 | $4.16 \times 10^{-1}$ |
| rs2488279 | 1.95 | $1.63 \times 10^{-1}$ | 0.15 | $7.00 \times 10^{-1}$ |
| rs2505513 | 14.00 | $1.83 \times 10^{-4}$ | 0.54 | $4.62 \times 10^{-1}$ |
| rs2505515 | 9.21 | $2.40 \times 10^{-3}$ | 3.65 | $5.61 \times 10^{-2}$ |
| rs2506024 | 13.11 | $2.93 \times 10^{-4}$ | 0.05 | $8.31 \times 10^{-1}$ |
| rs2796552 | 5.15 | $2.32 \times 10^{-2}$ | 0.71 | $3.99 \times 10^{-1}$ |
| rs2796575 | 0.81 | $3.68 \times 10^{-1}$ | 0.27 | $6.01 \times 10^{-1}$ |
| rs3004255 | 3.78 | $5.18 \times 10^{-2}$ | 0.83 | $3.61 \times 10^{-1}$ |
| rs3026693 | 0.51 | $4.75 \times 10^{-1}$ | 2.54 | $1.11 \times 10^{-1}$ |
| rs35092834 | 4.00 | $4.56 \times 10^{-2}$ | 0.25 | $6.19 \times 10^{-1}$ |
| rs3758514 | 2.71 | $9.96 \times 10^{-2}$ | 2.07 | $1.50 \times 10^{-1}$ |
| rs4422736 | 3.40 | $6.54 \times 10^{-2}$ | 0.00 | $9.92 \times 10^{-1}$ |
| rs4949062 | 2.26 | $1.33 \times 10^{-1}$ | 0.88 | $3.49 \times 10^{-1}$ |
| rs55737059 | 2.54 | $1.11 \times 10^{-1}$ | 0.65 | $4.21 \times 10^{-1}$ |
| rs57621833 | 2.90 | $8.85 \times 10^{-2}$ | 0.00 | $9.85 \times 10^{-1}$ |
| rs61844837 | 0.01 | $9.41 \times 10^{-1}$ | 0.38 | $5.36 \times 10^{-1}$ |
| rs6468008 | 5.88 | $1.53 \times 10^{-2}$ | 1.90 | $1.68 \times 10^{-1}$ |
| rs6974210 | 0.15 | $6.95 \times 10^{-1}$ | 0.06 | $7.99 \times 10^{-1}$ |
| rs7069590 | 2.59 | $1.07 \times 10^{-1}$ | 2.84 | $9.17 \times 10^{-2}$ |
| rs7074964 | 9.44 | $2.12 \times 10^{-3}$ | 1.09 | $2.97 \times 10^{-1}$ |
| rs72783255 | 1.06 | $3.02 \times 10^{-1}$ | 0.91 | $3.41 \times 10^{-1}$ |
| rs7393733 | 3.38 | $6.61 \times 10^{-2}$ | 0.41 | $5.22 \times 10^{-1}$ |
| rs7811476 | 0.05 | $8.26 \times 10^{-1}$ | 0.50 | $4.78 \times 10^{-1}$ |
| rs78146028 | 0.25 | $6.16 \times 10^{-1}$ | 0.49 | $4.84 \times 10^{-1}$ |
| rs7826312 | 0.74 | $3.91 \times 10^{-1}$ | 1.36 | $2.44 \times 10^{-1}$ |
| rs7835688 | 0.11 | $7.40 \times 10^{-1}$ | 0.17 | $6.84 \times 10^{-1}$ |
| rs788243 | 5.36 | $2.06 \times 10^{-2}$ | 3.29 | $6.97 \times 10^{-2}$ |
| rs788260 | 0.43 | $5.12 \times 10^{-1}$ | 1.93 | $1.65 \times 10^{-1}$ |
| rs788261 | 0.42 | $5.19 \times 10^{-1}$ | 1.93 | $1.65 \times 10^{-1}$ |
| rs788263 | 0.49 | $4.86 \times 10^{-1}$ | 1.93 | $1.65 \times 10^{-1}$ |
| rs788274 | 9.95 | $1.61 \times 10^{-3}$ | 1.44 | $2.30 \times 10^{-1}$ |
| rs869184 | 3.71 | $5.41 \times 10^{-2}$ | 0.08 | $7.81 \times 10^{-1}$ |
| rs953999 | 5.21 | $2.25 \times 10^{-2}$ | 5.24 | $2.21 \times 10^{-2}$ |

**Table S4:** Classification of AC-HSCR and HDRC probands by HSCR risk categories and risk scores.

| **Risk category** | **Sex** | **Segment Length** | **Familiality** | **Risk Score** | **Number (%) of AC-HSCR cases** | **Number (%) of HDRC cases** | **Combined number (%) of cases** |
| --- | --- | --- | --- | --- | --- | --- | --- |
| I | Male | Short | Simplex | 0 | 96 (35.7) | 80 (52.3) | 176 (41.7) |
| II | Male | Short | Multiplex | 1 | 31 (11.5) | 9 (5.9) | 40 (9.5) |
| III | Male | Long/TCA | Simplex | 1 | 44 (16.4) | 24 (15.7) | 68 (16.1) |
| IV | Male | Long/TCA | Multiplex | 2 | 29 (10.8) | 4 (2.6) | 33 (7.8) |
| V | Female | Short | Simplex | 1 | 23 (8.6) | 20 (13.1) | 43 (10.2) |
| VI | Female | Short | Multiplex | 2 | 12 (4.5) | 3 (2.0) | 15 (3.6) |
| VII | Female | Long/TCA | Simplex | 2 | 15 (5.6) | 7 (4.6) | 22 (5.2) |
| VIII | Female | Long/TCA | Multiplex | 3 | 19 (7.1) | 6 (3.9) | 25 (5.9) |

**Table S5:** Distribution of AC-HSCR and HDRC probands into HSCR risk score classes.

| **Cohort** | **Number (%) cases**<br>**with risk score 0** | **Number (%) of cases**<br>**with risk score 1** | **Number (%) of cases**<br>**with risk score 2/3** |
| --- | --- | --- | --- |
| **AC-HSCR** | 96 (35.7) | 98 (36.4) | 75 (27.9) |
| **HDRC** | 80 (52.3) | 53 (34.6) | 20 (13.1) |
| $\chi^2$ | 16.03 | | |
| ***P*** | 3.30×10<sup>-4</sup> |  |  |

Tests of heterogeneity ( $\chi^2$ ) and the significance of association ( $P$ ) are provided.

**Table S6:** Comparative summary of genetic associations at *RET*, *SEMA3* and *NRG1* variants in European ancestry HSCR subjects.

| ***RET*/rs2435357** |  |  |  |  |  |  |
| --- | --- | --- | --- | --- | --- | --- |
| Study | Sample size | Country | Variant | Risk allele count or frequency | Odds ratio | Ref. |
| Emison 2005 | 126 trios | USA | rs2435357 | T<sup>a</sup>:101; U<sup>b</sup>:25 | 4.0 | 11 |
| Emison 2010 | 519 trios | USA, France, Italy, Netherlands, Spain | rs2435357 | T<sub>hap</sub><sup>c</sup>: 0.57; U<sub>hap</sub><sup>d</sup>: 0.24 | 4.1<sup>i</sup> | 12 |
| Jiang 2015 | 649 trios | USA, France, Italy, Netherlands, Spain | rs2435357 | Case<sup>e</sup>: 0.61; Pseudo-control<sup>f</sup>: 0.21 | 5.3<sup>j</sup> | 1 |
| Kapoor 2015 | 355 cases; 633 controls | USA | rs2435357 | Case<sup>e</sup>: 0.58; Control<sup>g</sup>: 0.26 | 3.9 | 13 |
| Chatterjee 2016 | 346 cases; 732 controls | USA | rs2435357 | Case<sup>e</sup>: 0.58; Control<sup>g</sup>: 0.25 | 4.1 | 6 |
| Kapoor 2020 | 583 cases; ~7,500 controls | USA | rs2435357 | Case<sup>e</sup>: 0.60; Control<sup>g</sup>: 0.27 | 4.0 |  |
| Burzynski 2004 | 113 trios | Netherlands | rs2435362<sup>h</sup> | Case<sup>e</sup>: 0.68; Pseudo-control<sup>f</sup>: 0.26 | 6.0 | 14 |
| Vaclavikova 2014 | 162 cases; 205 controls | Czech Republic | rs2435357 | Case<sup>e</sup>: 0.72; Control<sup>g</sup>: 0.28 | 6.7 | 15 |
| Fadista 2018 | 170 cases; 4,717 controls | Denmark | rs2505994<sup>h</sup> | Case<sup>e</sup>: 0.63; Control<sup>g</sup>: 0.23 | 5.9 | 16 |
| Virtanen 2019 | 105 cases; 386 controls | Finland, Sweden | rs2435357 | Case<sup>e</sup>: 0.76; Control<sup>g</sup>: 0.32 | 5.7 | 17 |
| ***RET*/2506030** |  |  |  |  |  |  |
| Study | Sample size | Country | Variant | Risk allele count or frequency | Odds ratio | Ref. |
| Jiang 2015 | 649 trios | USA, France, Italy, Netherlands, Spain | rs2506030 | Case<sup>e</sup>: 0.58; Pseudo-control<sup>f</sup>: 0.41 | 2.0 | 1 |
| Kapoor 2015 | 355 cases; 633 controls | USA | rs2506030 | Case<sup>e</sup>: 0.56; Control<sup>g</sup>: 0.41 | 1.8 | 13 |
| Chatterjee 2016 | 346 cases; 732 controls | USA | rs2506030 | Case<sup>e</sup>: 0.56; Control<sup>g</sup>: 0.41 | 1.8 | 6 |
| Kapoor 2020 | 583 cases; ~7,500 controls | USA | rs2506030 | Case<sup>e</sup>: 0.56; Control<sup>g</sup>: 0.39 | 2.0 |  |
| Virtanen 2019 | 105 cases; 386 controls | Finland, Sweden | rs2506030 | Case<sup>e</sup>: 0.64; Control<sup>g</sup>: 0.41 | 2.3 | 17 |
| ***RET*/rs7069590** |  |  |  |  |  |  |
| Study | Sample size | Country | Variant | Risk allele count or frequency | Odds ratio | Ref. |
| Chatterjee 2016 | 346 cases; 732 controls | USA | rs7069590 | Case<sup>e</sup>: 0.84; Control<sup>g</sup>: 0.76 | 1.7 | 6 |
| Kapoor 2020 | 583 cases; ~7,500 controls | USA | rs7069590 | Case<sup>e</sup>: 0.84; Control<sup>g</sup>: 0.77 | 1.5 |  |
| ***SEMA3*/rs12707862** |  |  |  |  |  |  |
| Study | Sample size | Country | Variant | Risk allele count or frequency | Odds ratio | Ref. |
| Jiang 2015 | 649 trios | USA, France, Italy, Netherlands, Spain | rs12707682 | Case<sup>e</sup>: 0.35; Pseudo-control<sup>f</sup>: 0.23 | 1.8 | <sup>1</sup> |
| Kapoor 2015 | 355 cases; 633 controls | USA | rs12707682 | Case<sup>e</sup>: 0.30; Control<sup>g</sup>: 0.24 | 1.3 | <sup>13</sup> |
| Kapoor 2020 | 583 cases; ~7,500 controls | USA | rs12707682 | Case<sup>e</sup>: 0.31; Control<sup>g</sup>: 0.23 | 1.6 |  |
| Virtanen 2019 | 105 cases; 386 controls | Finland, Sweden | rs12707682 | Case<sup>e</sup>: 0.29; Control<sup>g</sup>: 0.19 | 1.7 | <sup>17</sup> |
| **SEMA3/rs11766001** |  |  |  |  |  |  |
| **Study** | **Sample size** | **Country** | **Variant** | **Risk allele count or frequency** | **Odds ratio** | **Ref.** |
| Jiang 2015 | 649 trios | USA, France, Italy, Netherlands, Spain | rs11766001 | Case<sup>e</sup>: 0.29; Pseudo-control<sup>f</sup>: 0.18 | 1.9<sup>k</sup> | <sup>1</sup> |
| Kapoor 2015 | 355 cases; 633 controls | USA | rs11766001 | Case<sup>e</sup>: 0.22; Control<sup>g</sup>: 0.15 | 1.6 | <sup>13</sup> |
| Kapoor 2020 | 583 cases; ~7,500 controls | USA | rs11766001 | Case<sup>e</sup>: 0.19; Control<sup>g</sup>: 0.14 | 1.5 |  |
| Virtanen 2019 | 105 cases; 386 controls | Finland, Sweden | rs11766001 | Case<sup>e</sup>: 0.22; Control<sup>g</sup>: 0.11 | 2.3 | <sup>17</sup> |
| **NRG1/rs16879552** |  |  |  |  |  |  |
| **Study** | **Sample size** | **Country** | **Variant** | **Risk allele count or frequency** | **Odds ratio** | **Ref.** |
| Kapoor 2015 | 355 cases; 633 controls | USA | rs16879552 | Case<sup>e</sup>: 0.03; Control<sup>g</sup>: 0.04 | 0.8 | <sup>13</sup> |
| Kapoor 2020 | 583 cases; ~7,500 controls | USA | rs16879552 | Case<sup>e</sup>: 0.05; Control<sup>g</sup>: 0.03 | 1.9 |  |

<sup>a</sup>Transmitted count; <sup>b</sup>Un-transmitted count; <sup>c</sup>Frequency in transmitted haplotypes; <sup>d</sup>Frequency in un-transmitted haplotypes; <sup>e</sup>Frequency in cases; <sup>f</sup>Frequency in pseudo-controls; <sup>g</sup>Frequency in controls; <sup>h</sup>in near-perfect LD with rs2435357; <sup>i</sup>Allele frequency shown is the average of allele frequencies reported from five cohorts and was used to calculate OR; <sup>j</sup>Allele frequency shown in the average of allele frequencies from primary GWAS and replication, which will lead to an OR of 5.9, however the OR shown is from the main text; <sup>k</sup>Allele frequency shown in the average of allele frequencies from primary GWAS and replication, and was used to calculate OR.

**Table S7:** Risk allele transmitted and un-transmitted counts in HSCR trios.

| Gene Locus | Variant | Coded allele | Transmitted/Un-transmitted counts | Transmission rate | *P* |
| --- | --- | --- | --- | --- | --- |
| *SEMA3* | rs10266471 | C | 89/86 | 0.51 | $8.21 \times 10^{-1}$ |
| *SEMA3* | rs1228877 | G | 146/134 | 0.52 | $4.73 \times 10^{-1}$ |
| *SEMA3* | rs55737059 | A | 75/54 | 0.58 | $6.45 \times 10^{-2}$ |
| *SEMA3* | rs1228937 | C | 126/107 | 0.54 | $2.13 \times 10^{-1}$ |
| *SEMA3* | rs11766001 | C | 102/75 | 0.58 | $4.24 \times 10^{-2}$ |
| *SEMA3* | rs2190050 | T | 92/111 | 0.45 | $1.82 \times 10^{-1}$ |
| *SEMA3* | rs35092834 | T | 139/102 | 0.58 | $1.72 \times 10^{-2}$ |
| *SEMA3* | rs12707682 | C | 138/119 | 0.54 | $2.36 \times 10^{-1}$ |
| *SEMA3* | rs7811476 | G | 136/138 | 0.50 | $9.04 \times 10^{-1}$ |
| *SEMA3* | rs6468008 | G | 152/142 | 0.52 | $5.60 \times 10^{-1}$ |
| *SEMA3* | rs17560655 | C | 99/79 | 0.56 | $1.34 \times 10^{-1}$ |
| *SEMA3* | rs6974210 | G | 69/64 | 0.52 | $6.65 \times 10^{-1}$ |
| *NRG1* | rs16879425 | C | 72/63 | 0.53 | $4.39 \times 10^{-1}$ |
| *NRG1* | rs10954845 | G | 114/101 | 0.53 | $3.75 \times 10^{-1}$ |
| *NRG1* | rs4422736 | T | 33/49 | 0.40 | $7.72 \times 10^{-2}$ |
| *NRG1* | rs7826312 | C | 149/110 | 0.58 | $1.54 \times 10^{-2}$ |
| *NRG1* | rs16879552 | T | 28/34 | 0.45 | $4.46 \times 10^{-1}$ |
| *NRG1* | rs7835688 | C | 174/133 | 0.57 | $1.93 \times 10^{-2}$ |
| *NRG1* | rs16879576 | A | 24/32 | 0.43 | $2.85 \times 10^{-1}$ |
| *RET* | rs11239767 | T | 90/91 | 0.50 | $9.41 \times 10^{-1}$ |
| *RET* | rs12570505 | T | 74/54 | 0.58 | $7.71 \times 10^{-2}$ |
| *RET* | rs3758514 | T | 168/132 | 0.56 | $3.77 \times 10^{-2}$ |
| *RET* | rs61844837 | C | 54/28 | 0.66 | $4.09 \times 10^{-3}$ |
| *RET* | rs2796552 | A | 131/108 | 0.55 | $1.37 \times 10^{-1}$ |
| *RET* | rs788243 | G | 198/130 | 0.60 | $1.74 \times 10^{-4}$ |
| *RET* | rs2796575 | A | 90/67 | 0.57 | $6.64 \times 10^{-2}$ |
| *RET* | rs788274 | A | 144/60 | 0.71 | $4.07 \times 10^{-9}$ |
| *RET* | rs788263 | G | 209/102 | 0.67 | $1.30 \times 10^{-9}$ |
| *RET* | rs788261 | C | 207/100 | 0.67 | $1.02 \times 10^{-9}$ |
| *RET* | rs788260 | A | 195/88 | 0.69 | $2.01 \times 10^{-10}$ |
| *RET* | rs3004255 | T | 184/130 | 0.59 | $2.31 \times 10^{-3}$ |
| *RET* | rs1935646 | A | 125/125 | 0.50 | $1.00 \times 10^0$ |
| *RET* | rs11819501 | C | 48/29 | 0.62 | $3.04 \times 10^{-2}$ |
| *RET* | rs2488279 | T | 144/106 | 0.58 | $1.63 \times 10^{-2}$ |
| *RET* | rs57621833 | T | 116/105 | 0.52 | $4.59 \times 10^{-1}$ |
| *RET* | rs1547930 | G | 128/79 | 0.62 | $6.60 \times 10^{-4}$ |
| *RET* | rs4949062 | C | 139/124 | 0.53 | $3.55 \times 10^{-1}$ |
| *RET* | rs2115816 | A | 160/147 | 0.52 | $4.58 \times 10^{-1}$ |
| *RET* | rs3026693 | A | 98/76 | 0.56 | $9.54 \times 10^{-2}$ |
| *RET* | rs7069590 | T | 132/68 | 0.66 | $6.03 \times 10^{-6}$ |
| *RET* | rs2435357 | T | 287/56 | 0.84 | $1.05 \times 10^{-35}$ |
| *RET* | rs12247456 | G | 165/68 | 0.71 | $2.09 \times 10^{-10}$ |
| *RET* | rs7393733 | G | 170/73 | 0.70 | $4.89 \times 10^{-10}$ |
| *RET* | rs2506024 | A | 199/65 | 0.75 | $1.62 \times 10^{-16}$ |
| *RET* | rs78146028 | A | 75/33 | 0.69 | $5.31 \times 10^{-5}$ |
| *RET* | rs12764797 | G | 75/27 | 0.74 | $2.01 \times 10^{-6}$ |
| *RET* | rs2505515 | G | 137/47 | 0.74 | $3.25 \times 10^{-11}$ |
| *RET* | rs1800860 | G | 130/102 | 0.56 | $6.60 \times 10^{-2}$ |
| *RET* | rs11238441 | C | 107/43 | 0.71 | $1.74 \times 10^{-7}$ |
| *RET* | rs2472737 | A | 136/97 | 0.58 | $1.06 \times 10^{-2}$ |
| *RET* | rs2505513 | C | 191/94 | 0.67 | $9.15 \times 10^{-9}$ |
| *RET* | rs10793423 | G | 185/133 | 0.58 | $3.55 \times 10^{-3}$ |
| *RET* | rs869184 | T | 216/104 | 0.68 | $3.83 \times 10^{-10}$ |
| *RET* | rs953999 | G | 169/80 | 0.68 | $1.70 \times 10^{-8}$ |
| *RET* | rs72783255 | C | 89/62 | 0.59 | $2.80 \times 10^{-2}$ |
| *RET* | rs7074964 | A | 173/106 | 0.62 | $6.04 \times 10^{-5}$ |

The tag variants at each locus are listed in genomic order. Transmitted and un-transmitted counts, transmission rate and significance of over-transmission (*P*) of coded allele (higher frequency in HSCR cases) at each variant are provided.

**Table S8:**
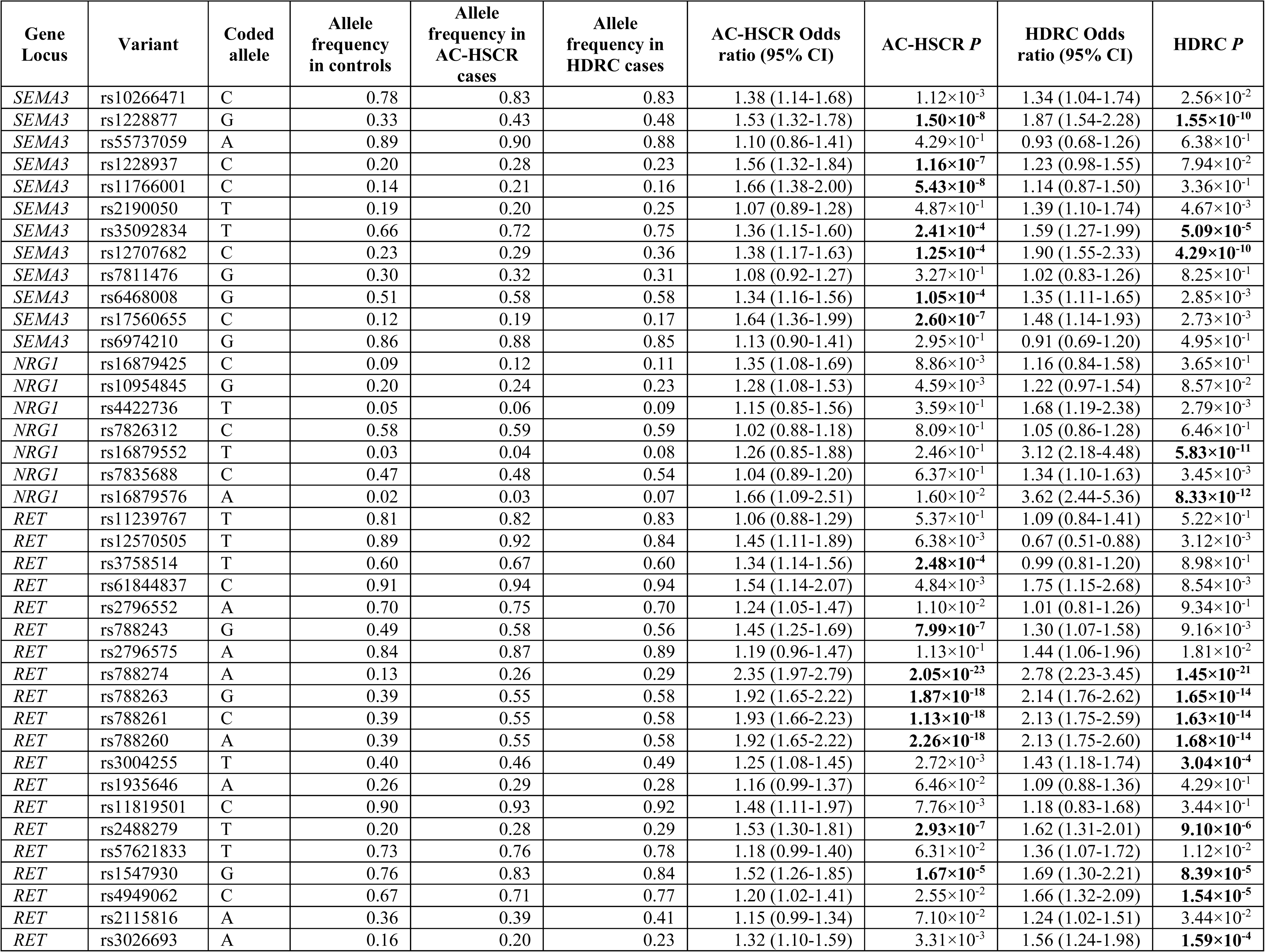
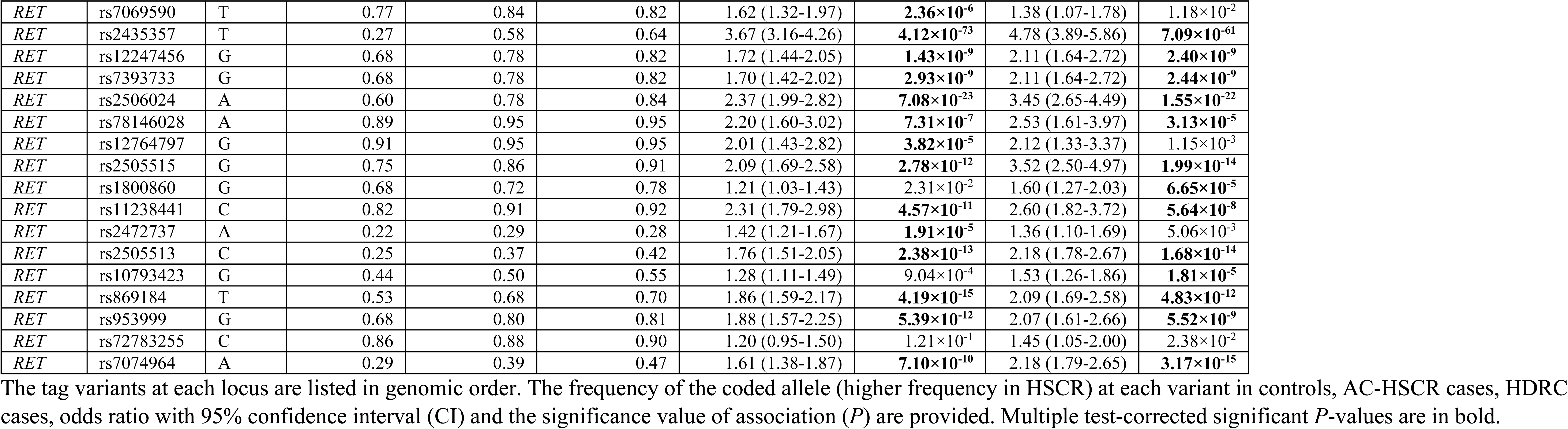
Case-control association tests for tag variants at *SEMA3*, *NRG1* and *RET* in AC-HSCR and HDRC Hirschsprung disease cohorts.

**Table S9:**
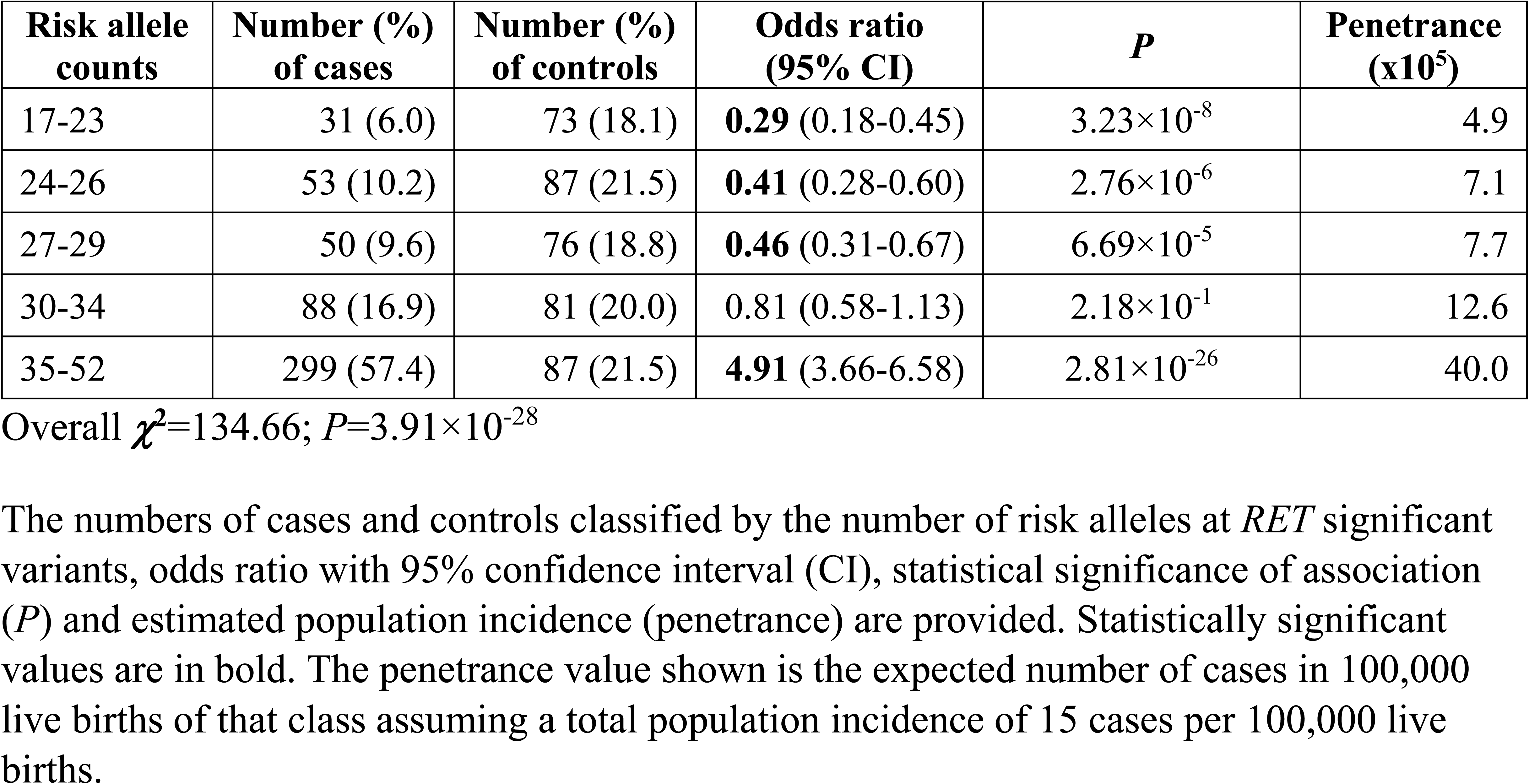
Odds ratio and risk of HSCR as a function of the number of risk-increasing variants at *RET*.

**Table S10:** Odds ratio and risk of HSCR as a function of the number of risk-increasing variants at *SEMA3*.

| Risk allele counts | Number (%) of cases | Number (%) of controls | Odds ratio (95% CI) | *P* | Penetrance (x10<sup>5</sup>) |
| --- | --- | --- | --- | --- | --- |
| 1-3 | 90 (16.4) | 92 (22.8) | **0.66** (0.48-0.92) | 0.014 | 10.8 |
| 4-5 | 117 (21.3) | 93 (23.0) | 0.91 (0.67-1.23) | 0.530 | 13.9 |
| 6-7 | 91 (16.6) | 75 (18.6) | 0.87 (0.62-1.22) | 0.424 | 13.4 |
| 8-9 | 125 (22.8) | 67 (16.6) | **1.48** (1.07-2.06) | 0.019 | 20.6 |
| 10-16 | 126 (23.0) | 77 (19.1) | 1.26 (0.92-1.74) | 0.148 | 18.1 |

Overall $\chi^2=11.87$ ; $P=0.018$
The numbers of cases and controls classified by the number of risk alleles at *SEMA3* significant variants, odds ratio with 95% confidence interval (CI), statistical significance of association (*P*) and estimated population incidence (penetrance) are provided. Statistically significant values are in bold. The penetrance value shown is the expected number of cases in 100,000 live births of that class assuming a total population incidence of 15 cases per 100,000 live births.

**Appendix S1:** List of HDRC investigators, sites and number of participant samples collected.

| **Investigators** | **Site & Location** | **Number of participant samples** |
| --- | --- | --- |
| Nicole Chandler (PI) and Hector Monforte (PI) | All Children's Hospital, St. Petersburg, FL | 17 |
| Philip Frykman (PI) | Cedars-Sinai Medical Center, Los Angeles, CA | 17 |
| Douglas Tamura (PI) | Children's Hospital Central California (Valley Children's Hospital), Madera, CA | 48 |
| Jason Frischer (PI) | Cincinnati Children's Hospital Medical Center, Cincinnati, OH | 244 |
| David A. Rodeberg (PI) | East Carolina University, Greenville, NC | 5 |
| Megan Durham (PI) | Emory/Children's Atlanta, Atlanta, GA | 45 |
| Jacob Langer (PI) | Hospital for Sick Children, Toronto, Canada | 49 |
| Isam Nasr (PI) | Johns Hopkins University, Baltimore, MD | 8 |
| Donald Shaul (PI) | Kaiser Permanente Southern California, Los Angeles, CA | 9 |
| Robert Cina (PI) | Medical University of South Carolina, Charleston, SC | 12 |
| Cheryl Gariepy (PI) | Nationwide Children's Hospital, Columbus, OH | 11 |
| Lisa McMahon (PI) | Phoenix Children's Hospital, Phoenix, AZ | 21 |
| Raj Kapur (PI) and Jeffrey Avansino (PI) | Seattle Children's Hospital, Seattle, WA | 33 |
| Monica Lopez (PI) | Texas Children's/Baylor College of Medicine, Houston, TX | 23 |
| Robert Russell (PI) | University of Alabama at Birmingham, Birmingham, AL | 6 |
| Daniel DeUgarte (PI) | University of California, Los Angeles, Los Angeles, CA | 21 |
| Samir K. Gadepalli (PI) | University of Michigan, Ann Arbor, MI | 5 |
| Michael Rollins (PI) | University of Utah, Salt Lake City, UT | 62 |
| Suyin Lum Min (PI) | Winnipeg Children's Hospital, Winnipeg, Canada | 12 |

## Notes

### Competing Interest Statement

The authors have declared no competing interest.

## References

1. Chakravarti A, Lyonnet S. Hirschsprung disease. In: Scriver CR, Beaudet AL, Valle D, et al, eds. The metabolic and molecular bases of inherited disease (8th edition). New York: McGraw- Hill; 2001:6231–6255.

2. Badner JA, Sieber WK, Garver KL, Chakravarti A. A genetic study of hirschsprung disease. Am J Hum Genet. 1990;46(3):568–580.

3. Parisi MA, Kapur RP. Genetics of hirschsprung disease. Curr Opin Pediatr. 2000;12(6):610–617.

4. Amiel J, Sproat-Emison E, Garcia-Barcelo M, et al. Hirschsprung disease, associated syndromes and genetics: A review. J Med Genet. 2008;45(1):1–14.

5. Alves MM, Sribudiani Y, Brouwer RW, et al. Contribution of rare and common variants determine complex diseases-hirschsprung disease as a model. Dev Biol. 2013;382(1):320–329.

6. Jiang Q, Arnold S, Heanue T, et al. Functional loss of semaphorin 3C and/or semaphorin 3D and their epistatic interaction with ret are critical to hirschsprung disease liability. Am J Hum Genet. 2015;96(4):581–596.

7. Emison ES, McCallion AS, Kashuk CS, et al. A common sex-dependent mutation in a RET enhancer underlies hirschsprung disease risk. Nature. 2005;434(7035):857–863.

8. Emison ES, Garcia-Barcelo M, Grice EA, et al. Differential contributions of rare and common, coding and noncoding ret mutations to multifactorial hirschsprung disease liability. Am J Hum Genet. 2010;87(1):60–74.

9. Garcia-Barcelo MM, Tang CS, Ngan ES, et al. Genome-wide association study identifies NRG1 as a susceptibility locus for hirschsprung’s disease. Proc Natl Acad Sci U S A. 2009;106(8):2694–2699.

10. Tilghman JM, Ling AY, Turner TN, et al. Molecular genetic anatomy and risk profile of hirschsprung’s disease. N Engl J Med. 2019;380(15):1421–1432. doi: 10.1056/NEJMoa1706594 [doi].

11. Kapoor A, Jiang Q, Chatterjee S, et al. Population variation in total genetic risk of hirschsprung disease from common RET, SEMA3 and NRG1 susceptibility polymorphisms. Hum Mol Genet. 2015;24(10):2997–3003.

12. Chatterjee S, Kapoor A, Akiyama JA, et al. Enhancer variants synergistically drive dysfunction of a gene regulatory network in hirschsprung disease. Cell. 2016;167(2):355–368.e10.

13. ENCODE Project Consortium, Bernstein BE, Birney E, et al. An integrated encyclopedia of DNA elements in the human genome. Nature. 2012;489(7414):57–74.

14. Roadmap Epigenomics Consortium, Kundaje A, Meuleman W, et al. Integrative analysis of 111 reference human epigenomes. Nature. 2015;518(7539):317–330.

15. Segal E, Raveh-Sadka T, Schroeder M, Unnerstall U, Gaul U. Predicting expression patterns from regulatory sequence in drosophila segmentation. Nature. 2008;451(7178):535–540.

16. International HapMap Consortium. A haplotype map of the human genome. Nature. 2005;437(7063):1299–1320.

17. 1000 Genomes Project Consortium. A map of human genome variation from population- scale sequencing. Nature. 2010;467(7319):1061–1073.

18. Bolk S, Pelet A, Hofstra RM, et al. A human model for multigenic inheritance: Phenotypic expression in hirschsprung disease requires both the RET gene and a new 9q31 locus. Proc Natl Acad Sci U S A. 2000;97(1):268–273. doi: 10.1073/pnas.97.1.268 [doi].

19. Gabriel SB, Salomon R, Pelet A, et al. Segregation at three loci explains familial and population risk in hirschsprung disease. Nat Genet. 2002;31(1):89–93. doi: 10.1038/ng868 [doi].

20. Kapoor A, Lee D, Zhu L, et al. Multiple SCN5A variant enhancers modulate its cardiac gene expression and the QT interval. Proc Natl Acad Sci U S A. 2019;116(22):10636–10645. doi: 10.1073/pnas.1808734116 [doi].

21. Tang CS, Tang WK, So MT, et al. Fine mapping of the NRG1 hirschsprung’s disease locus. PLoS One. 2011;6(1):e16181.

22. Rao SS, Huntley MH, Durand NC, et al. A 3D map of the human genome at kilobase resolution reveals principles of chromatin looping. Cell. 2014;159(7):1665–1680.

23. Barrett JC, Fry B, Maller J, Daly MJ. Haploview: Analysis and visualization of LD and haplotype maps. Bioinformatics. 2005;21(2):263–265.

24. Lek M, Karczewski KJ, Minikel EV, et al. Analysis of protein-coding genetic variation in 60,706 humans. Nature. 2016;536(7616):285–291.

25. Turner TN, Sharma K, Oh EC, et al. Loss of delta-catenin function in severe autism. Nature. 2015;520(7545):51–56. doi: 10.1038/nature14186 [doi].

26. Fadista J, Lund M, Skotte L, et al. Genome-wide association study of hirschsprung disease detects a novel low-frequency variant at the RET locus. Eur J Hum Genet. 2018;26(4):561–569. doi: 10.1038/s41431-017-0053-7 [doi].

27. Li Q, Zhang Z, Diao M, et al. Cumulative risk impact of RET, SEMA3, and NRG1 polymorphisms associated with hirschsprung disease in han chinese. J Pediatr Gastroenterol Nutr. 2017;64(3):385–390. doi: 10.1097/MPG.0000000000001263 [doi].

28. Jiang Q, Arnold S, Heanue T, et al. Functional loss of semaphorin 3C and/or semaphorin 3D and their epistatic interaction with ret are critical to hirschsprung disease liability. Am J Hum Genet. 2015;96(4):581–596.

29. International HapMap Consortium. A haplotype map of the human genome. Nature. 2005;437(7063):1299–1320.

30. 1000 Genomes Project Consortium. A map of human genome variation from population-scale sequencing. Nature. 2010;467(7319):1061-1073.

31. ENCODE Project Consortium, Bernstein BE, Birney E, et al. An integrated encyclopedia of DNA elements in the human genome. Nature. 2012;489(7414):57–74.

32. Roadmap Epigenomics Consortium, Kundaje A, Meuleman W, et al. Integrative analysis of 111 reference human epigenomes. Nature. 2015;518(7539):317–330.

33. Chatterjee S, Kapoor A, Akiyama JA, et al. Enhancer variants synergistically drive dysfunction of a gene regulatory network in hirschsprung disease. Cell. 2016;167(2):355–368.e10.

34. Lek M, Karczewski KJ, Minikel EV, et al. Analysis of protein-coding genetic variation in 60,706 humans. Nature. 2016;536(7616):285–291.

35. Barrett JC, Fry B, Maller J, Daly MJ. Haploview: Analysis and visualization of LD and haplotype maps. Bioinformatics. 2005;21(2):263–265.

36. Garcia-Barcelo MM, Tang CS, Ngan ES, et al. Genome-wide association study identifies NRG1 as a susceptibility locus for hirschsprung’s disease. Proc Natl Acad Sci U S A. 2009;106(8):2694–2699.

37. Pollard KS, Hubisz MJ, Rosenbloom KR, Siepel A. Detection of nonneutral substitution rates on mammalian phylogenies. Genome Res. 2010;20(1):110–121.

38. Emison ES, McCallion AS, Kashuk CS, et al. A common sex-dependent mutation in a RET enhancer underlies hirschsprung disease risk. Nature. 2005;434(7035):857–863.

39. Emison ES, Garcia-Barcelo M, Grice EA, et al. Differential contributions of rare and common, coding and noncoding ret mutations to multifactorial hirschsprung disease liability. Am J Hum Genet. 2010;87(1):60–74.

40. Kapoor A, Jiang Q, Chatterjee S, et al. Population variation in total genetic risk of hirschsprung disease from common RET, SEMA3 and NRG1 susceptibility polymorphisms. Hum Mol Genet. 2015;24(10):2997–3003.

41. Burzynski GM, Nolte IM, Osinga J, et al. Localizing a putative mutation as the major contributor to the development of sporadic hirschsprung disease to the RET genomic sequence between the promoter region and exon 2. Eur J Hum Genet. 2004;12(8):604–612. doi: 10.1038/sj.ejhg.5201199 [doi].

42. Vaclavikova E, Dvorakova S, Skaba R, et al. RET variants and haplotype analysis in a cohort of czech patients with hirschsprung disease. PLoS One. 2014;9(6):e98957. doi: 10.1371/journal.pone.0098957 [doi].

43. Fadista J, Lund M, Skotte L, et al. Genome-wide association study of hirschsprung disease detects a novel low-frequency variant at the RET locus. Eur J Hum Genet. 2018;26(4):561–569. doi: 10.1038/s41431-017-0053-7 [doi].

44. Virtanen VB, Salo PP, Cao J, et al. Noncoding RET variants explain the strong association with hirschsprung disease in patients without rare coding sequence variant. Eur J Med Genet. 2019;62(4):229–234. doi: S1769-7212(18)30076-4 [pii].

